# ACB1801 enhances tumor immunogenicity by targeting glycolysis/ferroptosis vulnerability and activating STAT1-signaling to overcome anti-PD-1 resistance in MSS colorectal cancer

**DOI:** 10.64898/2026.02.23.707369

**Authors:** Ruize Gao, Kris Van Moer, Coralie Pulido, Anaïs Oudin, Chenxi Li, Margaux Poussard, Teresa L. Ramos, Diane Murera, Elisabetta Bartolini, Annette Ives, Marine Gerbé de Thoré, Michele Mondini, Eric Deutsch, Guy Berchem, Christian Auclair, Bassam Janji

**Affiliations:** Tumor Immunotherapy and Microenvironment (TIME) group; Department of Cancer Research, Luxembourg Institute of Health, 6A, Rue Nicolas-Ernest Barblé, L-1210 Luxembourg City, Luxembourg; Animal Experimental Core Facility, Department of Cancer Research, Luxembourg Institute of Health, 29, rue Henri Koch, L-4354 Esch-sur-Alzette, Luxembourg; Faculty of Science, Technology and Medicine, University of Luxembourg, 2 place de l’Université, L-4365 Esch-sur-Alzette, Luxembourg; AC Bioscience SA. Biopôle Route de la Corniche 4 – Lysine. CH-1066 Epalinges, Switzerland; Gustave Roussy, Université Paris-Saclay, INSERM U1030, Villejuif, France; Department of Hemato-Oncology, Centre Hospitalier du Luxembourg, Luxembourg L-1210, Luxembourg; Department of Life Sciences and Medicine (DLSM), University of Luxembourg, Esch-sur-Alzette, Luxembourg; AC BioTech, Villejuif BioPark, Cancer Campus, F-94800 Villejuif, France

**Keywords:** Microsatellite stable colorectal cancer, harmine derivative ACB1801, immune checkpoint blockade, MHC-I antigen presentation, CXCL10, glycolysis, ferroptosis, tumor microenvironment

## Abstract

**Background:** Immune checkpoint blockade (ICB) therapies demonstrate low efficacy in microsatellite stable (MSS) colorectal cancer (CRC) due to an immune-desert tumor microenvironment (TME) characterized by low antigen presentation and limited tumor-infiltrating lymphocytes (TILs).

Harmine, a natural small-molecule and its promising derivatives ACB1801 have shown anti-tumor potential in preclinical models; however, their potential to reprogram the TME and overcome ICB resistance in MSS CRC remains unexplored. This study investigates whether and how ACB1801 can reshape TME to sensitize MSS CRC to ICB therapies.

**Methods:** We used the CT26 MSS colorectal cancer mouse model to evaluate the ability of the harmine derivative ACB1801 to enhance the efficacy of anti-PD-1 therapy. To characterize its mode of action, we performed immune landscape analysis and transcriptomic profiling of both CD45- and CD45+ tumor-derived cells. In parallel, mechanistic studies were conducted in vitro using mouse and human MSS CRC cell lines.

**Results:** We demonstrate that the harmine derivative ACB1801 enhances the effectiveness of anti-PD-1 therapy in an MSS CRC mouse model. Combination therapy significantly increased CD8+ T cell infiltration and reduced regulatory T-cell (Treg) density in the TME. Transcriptomic profiling of CRC cells isolated from tumors treated with either anti-PD-1 alone or in combination with ACB1801 revealed significant enrichment of metabolic pathways in the combination group, characterized by reduced glycolysis and enhanced ferroptosis signatures. These findings were supported by in vitro data showing that ACB1801 reduces tumor cell glycolytic activity and promotes ferroptotic vulnerability. Mechanistically, ACB1801 induced STAT1 signaling, promoted CXCL10 release, and enhanced major histocompatibility complex class I (MHC-I)-dependent antigen presentation on tumor cells, thereby increasing tumor susceptibility to anti-PD-1 therapy.

**Conclusion:** Collectively, our findings indicate that combination therapy with harmine derivatives and ICBs represents a promising strategy for treating MSS CRC patients.

## Background

Colorectal cancer (CRC) is the most common malignancy of the gastrointestinal tract and represents the third leading cause of cancer-related mortality worldwide [1]. Immunotherapy, particularly ICB targeting programmed cell death protein 1 (PD-1) and its ligand programmed Death-Ligand 1(PD-L1), has revolutionized the treatment of several cancers, including CRC with deficient mismatch repair (dMMR) and high microsatellite instability (MSI-H) [2]. However, the vast majority of CRC cases (approximately 80-85%) exhibit proficient mismatch repair (pMMR) and are either MSS or display low microsatellite instability (MSI-L). These tumors are typically considered immunologically “cold”, characterized by poor immune infiltration and resistance to ICB therapy [3, 4]. Therefore, therapeutic strategies designed to convert “cold” MSS CRC tumors into “hot,” immunoresponsive tumors hold great promise for broadening the clinical benefit of ICB in CRC patients.

The TME of MSS CRC is typically characterized by immune exclusion or an immune-desert phenotype, defined by low infiltration of TILs, a defect of CTLs, or both [5]. This immunologically “cold” state is driven by a combination of low TMB, impaired antigen presentation, and reduced surface expression of MHC-I molecules on APCs [6]. Together, these factors impair immune recognition and limit effective antitumor cytotoxicity. Therefore, therapeutic strategies that restore MHC-I expression and enhance antigen presentation represent a promising strategy to promote CTL infiltration and sensitize MSS CRC tumors to immune checkpoint blockade.

Chemokines are key regulators of the TME, orchestrating the recruitment and spatial distribution of immune cells [7]. In CRC, the chemokine network significantly influences both the extent and composition of immune cell infiltration, particularly cytotoxic T lymphocytes and other effector populations. Among these, CXCL10 is an interferon-inducible chemokine secreted by multiple cell types, including monocytes, endothelial cells, fibroblasts, and tumor cells. Through its interaction with the chemokine receptor CXCR3 on activated T cells, CXCL10 plays a pivotal role in anti-tumor immunity by promoting effector T cells trafficking into the TME [8]. By facilitating T cell infiltration and accumulation, CXCL10 contributes to the establishment of an inflamed, “hot” TME. Thus, strategies designed to boost CXCL10 expression within tumors may enhance effector T cell recruitment and improve the efficacy of immune checkpoint blockade.

Harmine is a naturally occurring β-carboline alkaloid first isolated from the seeds of *Peganum harmala* L. (Zygophyllaceae) [9]. It exhibits diverse biological and pharmacological activities, including anti-inflammatory, antiviral, antifungal, and antitumor effects [10], and has recently been suggested as a potential therapeutic agent for Alzheimer’s disease [11]. The antitumor properties of harmine have been investigated across multiple cancer types, where it modulates key cellular processes and signaling pathways, including epithelial-mesenchymal transition (EMT) [12], apoptosis [13], cell cycle progression [14-16], angiogenesis [17], and actin cytoskeleton remodeling [18]. In addition, harmine can overcome chemotherapeutic drug resistance by inhibiting of DYRK1A [19].

ACB1801 is a synthetic derivative of harmine, a naturally occurring β-carboline alkaloid found in *Peganum harmala*. Our previous study showed that ACB1801 enhances anti-PD-1 therapy in a B16-F10 melanoma syngeneic mouse model by upregulating MHC-I antigen presentation [20]. However, the potential of ACB1801 to improve anti-PD-1 therapy in MSS CRC and the underlying mechanisms remain largely unexplored.

In this study, we demonstrate that ACB1801 enhances the therapeutic efficacy of anti-PD-1 in a MSS CRC mouse model and reveals how it reshapes tumor cells and the tumor microenvironment. By modulating both the immune landscape and tumor cell metabolism, ACB1801 not only fosters immune cell infiltration and MHC-I expression on the tumors but also shifts cancer cells toward a state of metabolic vulnerability, sensitizing them to ferroptotic cell death.

## Methods

### Cell lines, cell culture, siRNA transfection, and reagents

CT26, MC38 and HT29 cell lines were obtained from American Type Culture Collection (ATCC). All cell lines used in this study were authenticated when applicable and routinely tested for mycoplasma contamination every 2 weeks. The cell lines were cultured as follows: CT26 with RPM-1640, MC38 with DMEM high-glucose, and HT29 in McCoy’s 5a Medium. All media were supplemented with 10% fetal calf serum (FCS; Sigma-Aldrich), 2⍰mM L-Glutamine (Sigma-Aldrich), 100⍰U/ml penicillin (Sigma-Aldrich), and 100⍰μg/ml streptomycin (Sigma-Aldrich). Cells were maintained in a humidified incubator at 37°C with 5% CO2.

siRNA transfections were performed using Lipofectamine RNAiMAX (Invitrogen, CAT# 13778150) according to the manufacturer’s instructions. Details of the siRNAs are provided in **Table 1**. The harmine derivative ACB1801 was kindly provided by AC Bioscience (Switzerland), and fer-1(ferrostatin-1) was purchased from Selleck chemicals (catalog No.S7243) to inhibit ferroptosis.

**Table 1.**
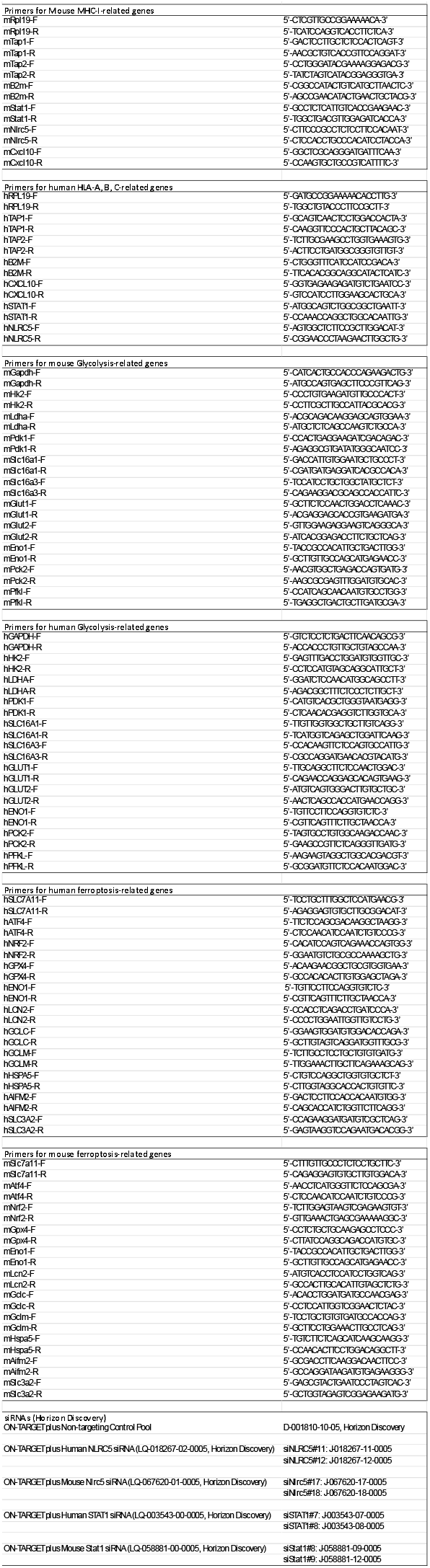
Summary of mouse and human primers, as well as siRNAs, used in this study.

### In vivo experiments

All animal experiments were conducted in compliance with European Union guidelines. The experimental protocols were approved by the Ethical Committee of the Luxembourg Institute of Health (LIH), the Animal Welfare Society, and the Luxembourg Ministry of Agriculture, Viticulture, and Rural Development (Approval No. LECR-2018-12).

Six-week-old BALB/c mice were purchased from Janvier Laboratories and housed in a certified animal facility under a 12-h light/dark cycle in a temperature-controlled room (22⍰± ⍰1°C), with free access to food and water, in accordance with Luxembourg regulations. For tumor transplantation, CT26 cells (0.5 × 10^6^ in 100⍰µL PBS) were implanted subcutaneously into the right flank of mice. Treatments were initiated as soon as the tumors became palpable. Mice received either vehicle solution (0.5% methylcellulose; Sigma, Cat# 9004-67-5) or ACB1801 (50⍰mg/kg daily via oral gavage). Isotype control antibody (BioXCell, Cat# BE0089, Clone 2A3) or anti-PD-1 antibody (CD279; BioXCell, Cat# BE0146, Clone RMP1-14) was administered intraperitoneally at 1⍰mg/kg every other day.

Tumor dimensions (length and width) were measured twice weekly using a caliper, and tumor volumes were calculated according to the formula: volume = length × width^2^ × 0.52. At the experimental endpoint, tumor tissues were collected for downstream analyses.

### Flow cytometry

For in vitro surface staining, cultured cells were washed with PBS and detached using 0.05% Trypsin-EDTA (Thermo Fisher Scientific, Cat# 25300062). Cells were quenched with a medium containing FBS, centrifuged at 400 × g for 5⍰min, and resuspended in MACS buffer (Miltenyi Biotec, Cat# 130-091-221) to remove cell debris. After a second centrifugation (400 × g, 5⍰min), the pellet was resuspended in MACS buffer, and cells were counted and adjusted to a minimum concentration of 1 × 10^7^ cells/ml. A total of 5 × 10^6^ cells per sample were incubated with the indicated antibodies at 4⍰°C for 301min. Samples not analyzed immediately were fixed with 2% PFA.

For immunophenotyping of tumor-infiltrating immune cells, tumors were dissociated using a tumor dissociation kit (Miltenyi Biotec, Cat# 130-096-730) according to the manufacturer’s instructions. Single-cell suspensions were resuspended in MACS buffer at 1 × 10^6^ cells/200⍰µl in U-bottom 96-well plates. After centrifugation (400 × g, 5⍰min), the cells were incubated on ice for 5⍰min with 10⍰µl of diluted anti-mouse CD16/32 antibody (BioLegend, Cat# 101319, 1:10 dilution) to block Fc receptors, followed by incubation with antibody cocktails (30⍰µl/well) for 30⍰min at 4⍰°C. The cells were then washed and resuspended in MACS buffer. Samples not requiring FoxP3 staining were fixed with 2% PFA, while those requiring FoxP3 analysis were processed according to the manufacturer’s protocol for the eBioscience Foxp3/Transcription Factor Staining Buffer Set (Thermo Fisher Scientific, Cat# 00-5523-00). Flow cytometry was performed on a NovoCyte Quanteon Flow Cytometer System (4 lasers). Data were analyzed using FlowJo software (FlowJo LLC, Ashland, OR).

The following antibodies from BioLegend were used in this study: PE anti-mouse H-2Kb (Cat# 116507, Clone AF6-88.5), PE anti-mouse H-2Kb-SIINFEKL (Cat# 141603, Clone 25-d1.16), APC anti-mouse H-2Kd (Cat# 116619, Clone SF1-1.1), PE anti-human HLA-A,B,C (Cat# 311405, Clone W6/32), Brilliant Violet (BV) 421 anti-mouse FoxP3 (Cat# 126419, Clone MF-14), BV570 anti-mouse CD44 (Cat# 103037, Clone IM7), BV605 anti-mouse CD69 (Cat# 104529, Clone H1.2F3), BV650 anti-mouse CD4 (Cat# 100469, Clone GK1.5), BV785 anti-mouse CD3 (Cat# 100231, Clone 17A2), APC anti-mouse CD8a (Cat# 100711, Clone 53-6.7), PE anti-mouse PD-1 (CD279; Cat# 135205, Clone 29F.1A12), PE/Cy5 anti-mouse CD25 (Cat# 102010, Clone PC61), PerCP/Cy5.5 anti-mouse TCRβ (Cat# 109227, Clone H57-597), PE/Cy7 anti-mouse CD19 (Cat# 152417, Clone 1D3/CD19), APC anti-mouse NK1.1 (Cat# 156505, Clone S17016D), BV421 anti-mouse Ly-6C (Cat# 128031, Clone HK1.4), BV510 anti-mouse I-A/I-E (Cat# 107635, Clone M5/114.15.2), BV605 anti-mouse F4/80 (Cat# 123133, Clone BM8), BV785 anti-mouse CD11b (Cat# 101243, Clone M1/70), PE anti-mouse CD206 (MMR; Cat# 141705, Clone C068C2), PE/Cy5 anti-mouse CD11c (Cat# 117316, Clone N418), and PE/Cy7 anti-mouse Ly-6G (Cat# 127617, Clone 1A8). The viability dye Zombie NIR (Cat# 423105) was included for assessing live/dead cells.

### *In vitro* OVA257-264(SIINFEKL) antigen presentation assay

MC38 cells were seeded at a density of 0.3 × 10^6^ cells per well in 6-well plates. After 24⍰h, the cells were incubated with 1⍰nmol OVA257–264 (SIINFEKL) peptide (InvivoGen, Cat# vac-sin) for 8⍰h, followed by treatment with either DMSO or ACB1801 for an additional 24⍰h. the cells were then harvested, and the surface expression of OVA-bound MHC-I was analyzed by flow cytometry as described in the “Flow cytometry” section.

### CD8+ OT-I T cell isolation and cytotoxicity assay

CD8+ OT-I T cells were isolated from the spleens/lymph nodes of OT-I mice (C57BL/6-Tg(TcraTcrb)100Mjb/Crl; Charles River Laboratories, France) using the MojoSort Mouse CD8 T Cell Isolation Kit (BioLegend, Cat# 4800008), according to the manufacturer’s instructions. Isolated CD8+ OT-I T cells were cultured in complete RPMI-1640 medium supplemented with 10% FBS, 20⍰mM HEPES, 2⍰mM L-glutamine, 1⍰mM sodium pyruvate, 1% penicillin-streptomycin, and 50⍰µM 2-mercaptoethanol. For long-term storage, cells were cryopreserved in 90% FBS and 10% DMSO, transferred into cryovials, cooled gradually in a Mr. Frosty freezing container at -80⍰°C, and subsequently transferred to liquid nitrogen for long-term storage. For activation, CD8+ OT-I T cells were stimulated with Dynabeads Mouse T-Activator CD3/CD28 (Thermo Fisher Scientific, Cat# 11452D) at a 1:1 bead-to-cell ratio. After 2 days, the beads were removed, and recombinant mouse IL-2 (20⍰ng/ml; PeproTech, Cat# 212-12-20UG) was added to the culture medium.

For the cytotoxicity assay, MC38 cells (5 × 10^3^ per well) were seeded into 96-well plates and pulsed with 1⍰nmol OVA257–264 (SIINFEKL) peptide (InvivoGen, Cat# vac-sin) for 8⍰h prior to co-culture. Activated CD8+ OT-I T cells were then added at different effector-to-target (E:T) ratios (0:1, 1:2.5, 1:1, and 2.5:1) and co-cultured for 48⍰h. Tumor cell viability was assessed by flow cytometry using Precision Count Beads (BioLegend, Cat# 424902).

### RNA extraction and quantitative RT-PCR

Total RNA was extracted using an RNA extraction kit (MACHEREY-NAGEL, Cat# 740984.250) according to the manufacturer’s instructions. First-strand cDNA synthesis was performed using the Maxima First Strand cDNA Synthesis Kit. Quantitative real-time PCR was carried out with SYBR qPCR Blue Master Mix (Eurogentec, Cat# UF-LSMT-B0701) on a StepOne Plus Real-Time PCR System (Applied Biosystems). Relative gene expression levels were calculated using the comparative Ct (ΔΔCt) method. Primer sequences are listed in **Table 1**.

### Cytokine/chemokine profiling

CT26 cells were seeded at 1 × 10^6^ cells per 60⍰mm dish. After 24⍰h, the cells were treated with either DMSO or ACB1801 for an additional 24⍰h. Supernatants were collected, centrifuged at 1,500⍰rpm for 10⍰min to remove cell debris, and retained for proteome profiler array analysis. The cells were harvested in RNA lysis buffer for qRT-PCR to assess the expression of the corresponding genes. Protein concentrations of the supernatants were measured using the Pierce BCA Protein Assay Kit (Thermo Fisher Scientific, Cat# 23227) and used to normalize proteome profiler array results. Proteome profiling was performed using a Mouse XL Cytokine Array Kit (Bio-Techne, Cat# ARY028) following the manufacturer’s protocol. All membranes were exposed to the same X-ray films with multiple exposure times to ensure optimal detection. The membranes were scanned using a high-resolution scanner, and dot intensities were quantified using ImageJ software.

### ELISA

Supernatants and mRNA were prepared and collected as described in the “Cytokine/chemokine profiling” section. Secreted CXCL10 levels were measured using a Human CXCL10/IP-10 DuoSet ELISA Kit (Bio-Techne, Cat# DY266) for HT29 cells and a Mouse CXCL10/IP-10/CRG-2 DuoSet ELISA Kit (Bio-Techne, Cat# DY466) for CT26 cells, following the manufacturer’s instructions.

### Tumor plasma collection

Tumors were harvested from mice and placed in 10⍰mL of serum-free medium in 50⍰mL tubes. The tumors were mechanically minced into small fragments and enzymatically digested using a tumor dissociation kit (Miltenyi Biotec, Cat# 130-096-730) following the manufacturer’s instructions. Single-cell suspensions were prepared by centrifugation at 1,500⍰rpm for 10⍰min at 4⍰°C. Supernatants were collected and either used immediately or frozen at -80⍰°C. To concentrate tumor plasma, the supernatants were centrifuged again at 1,500⍰rpm for 10⍰min at 4⍰°C to remove residual cell pellets. Ten milliliters of supernatant were loaded into protein concentration tubes (3K MWCO, Thermo Fisher Scientific, Cat# 88526) and ultracentrifuged at 8,000 × g for 2–5⍰h to reduce the volume from 10⍰mL to approximately 1⍰mL. Concentrated tumor plasma was aliquoted for downstream analyses or stored at -80⍰°C.

### Immunoblotting analysis

Cells were washed twice with 1× PBS and lysed in RIPA buffer (Sigma, Cat# R0278). Lysates were centrifuged to remove cell debris, and protein concentrations were determined prior to immunoblotting. Proteins were separated by SDS–PAGE and transferred to PVDF membranes. Membranes were blocked with 5% skimmed milk in Tris-buffered saline with Tween 20 (TBST) and incubated overnight at 4⍰°C with primary antibodies. After three 10-minute washes with TBST, the membranes were incubated with appropriate secondary antibodies for 2⍰h at room temperature, followed by three additional 10-minute washes. Chemiluminescent signals were detected using HRP substrate (Millipore, WBKLS0500) and X-ray films (FUJIFILM). The following antibodies were used: Anti-Stat1 (Cell Signaling Technology, Cat. #9172); Anti-Phospho Tyr701-Stat1 (Cell Signaling Technology 58D6 Cat. #9167); anti-beta-actin monoclonal antibody (Invitrogen BA3R Cat. #MA5-15739-HRP).

### Intracellular glucose measurement

Cells were seeded at 2 × 10^4^ cells per well in a 96-well plate. After 24⍰h, the cells were treated with DMSO or ACB1801 for an additional 24⍰h. Intracellular glucose levels were measured using the Glucose-Glo Assay kit (Promega, Cat# J6021) according to the manufacturer’s instructions. Briefly, the culture medium was removed, and cells were washed twice with 200⍰µL PBS per wash. Washed cells were resuspended in 25⍰µL PBS, followed by the addition of 12.5⍰µL inactivation solution and mixing on an orbital shaker for 5⍰min. Next, 12.5⍰µL of neutralization solution was added and mixed for 60⍰s. Fifty microliters of glucose detection reagent were added, and mixed for 30–60⍰s, and the plate was incubated for 60⍰min at room temperature. Luminescence was recorded, and glucose concentrations were calculated using a standard curve.

### Lactate measurement

Both extracellular and intracellular lactate levels were measured using the Lactate-Glo Assay Kit (Promega, Cat# J5021) according to the manufacturer’s instructions. Cells were seeded at 2 × 10^4^ cells per well in 96-well plates. After 24⍰h, the cells were treated with DMSO or ACB1801 for an additional 24⍰h, and samples were then collected. For extracellular lactate measurement, 5⍰µL of culture medium was collected and diluted in 95⍰µL PBS. Samples were kept at -20⍰°C until measurement. Fifty microliters of each sample were transferred to a 96-well plate, and 50⍰µL of lactate detection reagent was added to each well. Plates were shaken for 30–60⍰s, incubated for 60 min at room temperature, and luminescence was recorded. For intracellular lactate measurement, the culture medium was removed, and the cells were washed twice with 200⍰µL PBS per wash. The cells were resuspended in 25⍰µL PBS, followed by 12.5⍰µL inactivation solution, and mixed on an orbital shaker for 5⍰min. Next, 12.5⍰µL of neutralization solution was added and mixed for 60⍰s. Fifty microliters of lactate detection reagent were added, mixed for 30–60⍰s, and incubated for 60⍰min at room temperature. Luminescence was recorded, and lactate concentrations were calculated using a standard curve.

### GSH/GSSG ratio measurement

A GSH/GSSG-Glo Assay Kit (Promega, Cat# V6611) was used to measure reduced (GSH) and oxidized (GSSG) glutathione levels according to the manufacturer’s instructions. Two parallel 96-well plates were prepared under identical conditions. Cells were seeded at 5 × 10^3^ cells per well, and after 24⍰h, treated with DMSO or ACB1801 for an additional 24⍰h. Freshly prepared total glutathione lysis reagent and oxidized glutathione lysis reagent were added to each well (50⍰µL per well) on the respective plates, and the plates were shaken for 5⍰min at room temperature. Fifty microliters of luciferin generation reagent were then added to all wells, mixed briefly, and incubated at room temperature for 30⍰min. One hundred microliters of luciferin detection reagent were added, the plates were mixed briefly, incubated for 15⍰min, and luminescence was recorded. The GSH/GSSG ratio for vehicle-control cells was determined by subtracting the luminescence signal corresponding to oxidized glutathione (GSSG) from the total glutathione signal and then dividing this difference by half of the GSSG signal. The GSH/GSSG ratio for treated cells was calculated in the same manner, using the corresponding luminescence signals from treated wells.

### Cellular ROS measurement

Cells were seeded at 1 × 10^6^ per 60⍰mm dish. After 24⍰h, the cells were treated with DMSO or ACB1801 for an additional 24⍰h. For general ROS detection, the cells were stained with CellROX™ Green (Thermo Fisher Scientific, Cat# C10492) for 30⍰min at 37⍰°C, harvested by trypsinization, resuspended in PBS, and filtered through a 40⍰µm strainer. Flow cytometry analysis was performed using the NovoCyte Quanteon Flow Cytometer (4 lasers) with a 488⍰nm laser, and data were processed using FlowJo (FlowJo LLC, Ashland, OR). For lipid peroxidation measurement, cells were stained with C11-BODIPY 581/591 (Thermo Fisher Scientific, Cat# D3861) for 30⍰min at 37⍰°C, harvested, resuspended in PBS, and filtered through a 40⍰µm strainer. Flow cytometry was performed using 488⍰nm and 561⍰nm lasers, and data were analyzed in FlowJo.

### Cell viability assay

Cells were seeded at 5 × 10^3^ per well in a 96-well plate. After 24⍰h, the cells were treated with DMSO, ACB1801, or ACB1801 in combination with Ferrostatin-1 for an additional 24⍰h. The plates and CellTiter-Glo 2.0 Assay Solution (Promega, Cat# G9241) were equilibrated to room temperature. An equal volume of medium was added to each well, and the plates were mixed on an orbital shaker for approximately 2⍰min to induce cell lysis. The plates were then incubated at room temperature for 10⍰min to stabilize the luminescent signal, which was subsequently recorded.

### RNA-sequencing analysis

RNA was extracted in biological triplicates using the MACHEREY-NAGEL RNA Extraction Kit (Cat# 740984.250). RNA quality was assessed with a fragment analyzer (Labgene, DNF-471-0500 or DNF-472-0500), and concentrations were measured using the Quanti-iT™ RiboGreen RNA Assay Kit (Thermo Fisher Scientific). Libraries were prepared from 200⍰ng of RNA using the TruSeq Stranded Total RNA LT Sample Prep Kit (Illumina), and poly-A+ RNA was sequenced on an Illumina HiSeq 2500 using the HiSeq SBS Kit v4. Reads (81⍰bp, single-end) were aligned to the human genome (hg19, GRCh37.75) using RNA-STAR with default parameters, retaining only uniquely mapping reads and filtering reads without evidence in the spliced junction table. Gene-level expression counts were quantified over exons (RefSeq mRNA, UCSC, downloaded December 2015) using the qCount function from the QuasR package (v1.12.0). Differential expression analysis was performed with edgeR (v3.14.0), considering genes with p < 0.05 and |log2 fold change| ≥ 0.5 as significant. Selected genes were subjected to functional and pathway enrichment analysis.

### Functional enrichment analysis

Differentially expressed genes were analyzed for enrichment in biological processes and pathways using Bioconductor packages in R (org.Mm.eg.db v3.3.0, GO.db v3.4.1, GOstats v2.42.0, KEGG.db v3.2.3, ReactomePA v1.16.2). Significance was assessed using the hypergeometric test (Fisher’s exact test), with P ≤ 0.05 considered significant.

### Gene set enrichment analysis (GSEA)

GSEA was performed using a Java application from the Broad Institute (v3.0, http://www.broadinstitute.org/gsea). Gene sets were derived from gene ontology (GO) annotations, and pathway information was obtained from the Kyoto Encyclopedia of Genes and Genomes (KEGG, http://www.genome.jp/kegg/).

### Re-analysis of transcriptomic profiling data

RNA-sequencing gene expression data were retrieved from the TCGA COAD dataset (Cancer Genome Atlas Research Network, 2017) via cBioPortal (http://www.cbioportal.org).

### Statistical analysis

All statistical tests were two-sided. Data are presented as means, with error bars representing ± standard error of the mean (SEM). Analyses were performed using GraphPad Prism 9.0.

### Large Language Models (LLMs)

During the preparation of this grant, we used LLMs, including ChatGPT and Perplexity, to assist with summarizing content and improving the clarity and readability of the text. All generated material was carefully reviewed, critically evaluated, and edited as necessary by the authors.

## Results

### ACB1801 enhances the efficacy of anti-PD-1 therapy in the MSS CRC CT26 mouse model

We evaluated the effects of ACB1801 in combination with anti-PD-1 therapy on the MSS CRC CT26 syngeneic mouse model. Mice were treated with isotype control, ACB1801, anti-PD-1, or a combination of ACB1801 and anti-PD-1 **(Figure 1A)**. Tumor growth curves revealed that monotherapies with either ACB1801 or anti-PD-1 had no significant effects on tumor growth. Remarkably, the co-administration of ACB1801 with anti-PD-1 resulted in a significant reduction in both tumor volumes and tumor weights compared to anti-PD-1 alone (**Figure 1B and 1C**), translating into a marked improvement in overall survival (**Figure 1D**). Our data indicate that the harmine derivative ACB1801 enhances the efficacy of anti-PD-1 therapy in the MSS CRC mouse model.

**Figure 1.**
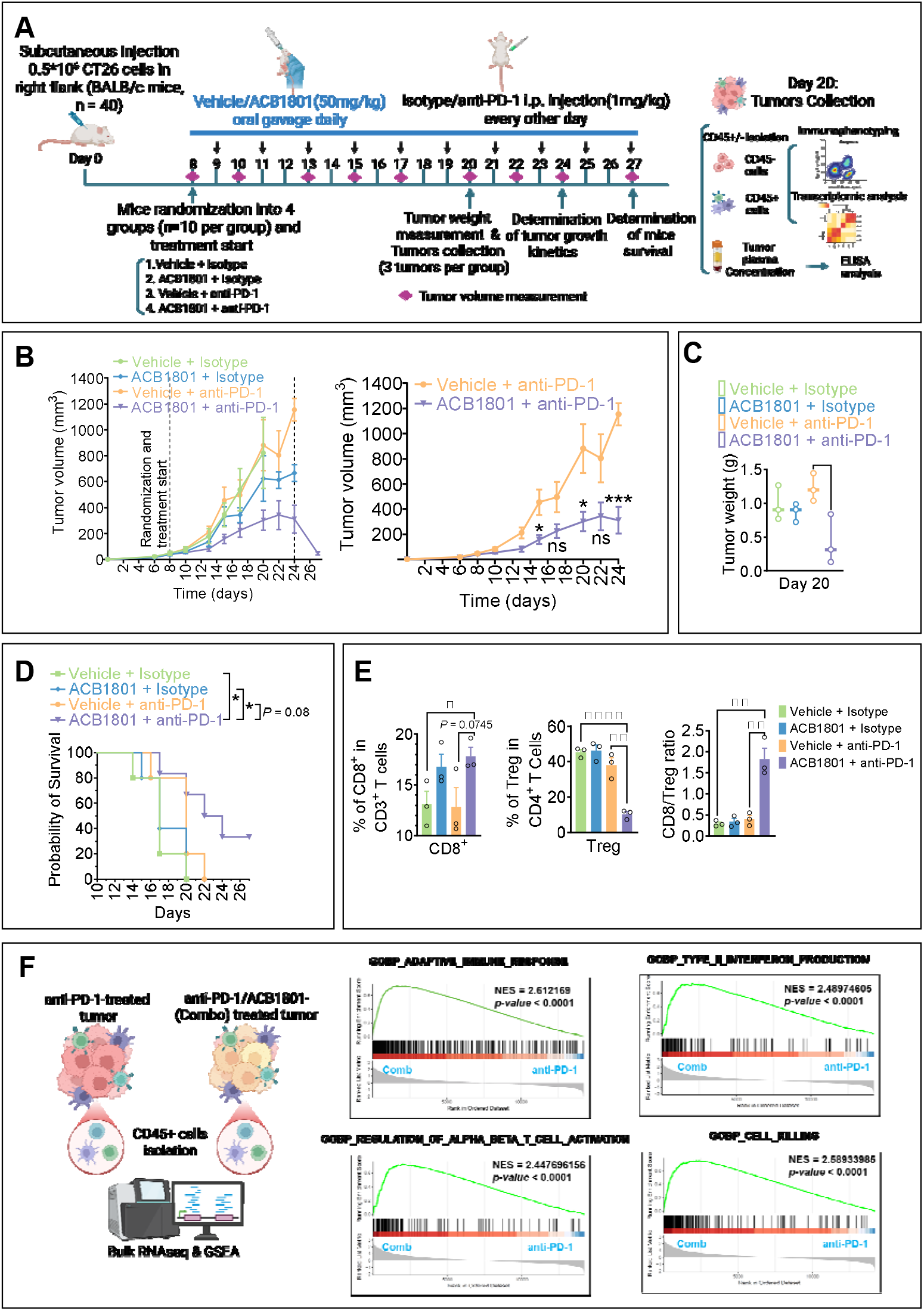
ACB1801 enhances anti-PD-1 efficacy in the MSS CRC CT26 syngeneic mouse model. **(A)** In vivo experimental design. CT26 cells (0.5 × 10^6^ cells per mouse) were injected subcutaneously into the right flank of 40 BALB/c mice. On day 8, mice were randomized into 4 treatment groups (n = 10 per group): (1) vehicle + isotype control, (2) ACB1801 + isotype control, (3) vehicle + anti–PD-1, and (4) ACB1801 + anti–PD-1, and treatments were initiated. Vehicle or ACB1801 (50 mg/kg) was administered daily by oral gavage, while isotype control or anti–PD-1 antibody (1 mg/kg) was administered intraperitoneally (i.p.) every other day. On day 20, group 1 was discontinued because 60% of mice reached humane endpoints or exhibited tumor necrosis or health deterioration. On the same day, three tumors per group were harvested, weighed, and dissociated for isolation of CD45- and CD45+ cells for transcriptomic analysis and immune phenotyping; corresponding tumor plasma samples were collected for cytokine and chemokine profiling. Tumor growth kinetics were assessed up to day 24, and overall survival was monitored until day 27, when the remaining mice were sacrificed. **(B)** Tumor growth kinetics of CT26 tumors in BALB/c mice treated with ACB1801 and/or anti–PD-1. Tumor volumes were monitored longitudinally and are presented as mean ± SEM. Upper panel represent the tumor growth kinetic of the four group plotted until day 27. Lower panels represent the comparison of: vehicle + isotype control versus ACB1801 + isotype control (left panel); vehicle + isotype control (group 1) versus vehicle + anti–PD-1 (middle panel); vehicle + anti–PD-1 versus the combination of ACB1801 and anti–PD-1 (right panel) plotted until day 24. Statistical significance difference in the tumor volume was determined using on day 24 was determined using unpaired two-tailed t test with Welch’s correction (ns = not significant; *p < 0.05; **p < 0.005). **(C)** Tumor weights on day 20 in each treatment group. Data represent the mean ± SEM of three tumors per group. Statistical significance was assessed using an unpaired two-tailed t-test with Welch’s correction (*p < 0.05). Only statistically significant differences are shown. **(D)** Kaplan–Meier survival curves of CT26 tumor–bearing mice. Loss of survival was defined as death, extensive tumor necrosis, significant body weight loss, deterioration of health status, or tumor volume exceeding 1500 mm^3^. Survival percentages were calculated using GraphPad Prism, and statistical significance was determined using the log-rank (Mantel–Cox) test (*p < 0.05). **(E)** Flow cytometry analysis of tumor-infiltrating immune cells. Left: CD8+ T cells; middle: Tregs; right: CD8/Treg ratio. Tumors were harvested on day 20, and immune populations were gated on live CD45+ cells. Each dot represents an individual tumor (n = 3 per group). Data are presented as the mean of 3 experiments ± SEM. Statistical significance was assessed using ordinary one-way ANOVA (*p < 0.05; **p < 0.001; ****p <0.00001). **(F)** Gene set enrichment analysis (GSEA) of RNA-seq data from isolated intratumoral CD45+ cells comparing the combination (combo) treatment group (ACB1801 + anti-PD-1) with the single-agent anti-PD-1 group. The enrichment plots show significant upregulation of gene sets associated with adaptive immune response, type II interferon production, regulation of alpha/beta T cell activation, cell killing in the combination group. For each pathway, the green line represents the running enrichment score, vertical black bars indicate the positions of genes from the gene set in the ranked list, and the normalized enrichment score (NES) and p-value are reported within each panel.

We next investigated whether the enhanced therapeutic benefit of anti-PD-1 therapy with ACB1801 was associated with changes in the immune landscape of the tumors. To explore this, comprehensive immune phenotyping was performed on tumors harvested from mice treated with the combination therapy, as well as those from the control and mono-therapy groups. Flow cytometry analysis revealed that tumors treated with the combination therapy showed increased CD8+ T cell infiltration, accompanied by a significant reduction in Tregs. Therefore, the CD8/Treg ratio was elevated in tumors treated with combination therapy compared to the control and single-treatment groups (**Figure 1E**). The combination therapy did not significantly alter the frequencies of total and activated (CD69+) CD4+ effector T cells, NK cells, CD11b+ myeloid cells, F4/80+ macrophages, PMN-MDSCs, monocytes, or CD11c+ dendritic cells (**Supplementary Figure 1A**). Interestingly, however, the proportion of PD-1+ CD4+ effector T cells was reduced in tumors treated with the combination therapy (**Supplementary Figure 1B**), suggesting a restoration of CD4+ T-cell functionality and alleviation of exhaustion. This finding is consistent with the enhanced antitumor immunity observed in this group of tumors.

Transcriptomic analysis of CD45+ immune cells isolated from tumors treated with anti-PD-1 or a combination of anti-PD-1 and ACB1801 showed that pathways related to adaptive immunity, type II interferon signaling, T cell activation, and cytotoxicity were significantly enriched in the combination therapy group versus the group treated with anti-PD-1 alone (**Figure 1F**), supporting a shift from an immune-suppressive to an immune-supportive TME. Moreover, enrichment of gamma-delta-T-cell activation indicates enhanced innate immune responses upon combination therapy (**Supplementary Figures 2A**). Pathways associated with cytokine production (**Supplementary Figures 2A**), along with elevated intratumoral IFNg levels (**Supplementary Figures 2B**) further confirm the activation of adaptive immunity. Interestingly, antigen-processing and presentation pathways were also enriched in the combination group, suggesting increased tumor infiltration of antigen-presenting cells (APCs), including dendritic cells, as well as maturation of M1-like tumor-associated macrophages (**Supplementary Figures 2A**).

### ACB1801 downregulates glycolysis and induces ferroptotic vulnerability in CRC tumor cells

Cancer cells undergo metabolic reprogramming to support survival, therapy resistance, and immune evasion [21, 22]. A dimeric form of β-carboline has been reported to impact sarcoma cell metabolism [23], but the underlying mechanisms remain incompletely understood. To investigate whether and how the β-carboline derivative ACB1801 affects tumor cells in vivo, we performed comparative transcriptomic analysis of CD45^−^ cells isolated from tumors treated with combination therapy (ACB1801 + anti-PD-1) versus those treated with anti-PD-1 monotherapy.

KEGG enrichment analysis revealed that among the top 30 enriched cellular functions, protein processing in the endoplasmic reticulum and ubiquitin-mediated proteolysis pathways were significantly represented in combination-treated tumors. These findings support the role of ACB1801 in enhancing MHC-I antigen processing and presentation. Remarkably, metabolic pathways were also significantly enriched in CD45-cells isolated from combination-treated tumors compared with those treated with anti-PD-1 monotherapy (**Figure 2A**). Detailed analysis of KEGG pathway annotations revealed that this metabolic reprogramming was characterized by an enrichment in carbohydrate, lipid, and amino acid metabolism, as well as glycan biosynthesis and metabolism (**Figure 2B**). These findings support the hypothesis that ACB1801 drives metabolic remodeling in tumor cells, which may contribute to improved anti-tumor responses. To further delineate the role of ACB1801 at the cellular level, we performed bulk RNA-sequencing on untreated or ACB1801-treated CT26 cells. Gene set enrichment analysis (GSEA) revealed a downregulation of glycolysis along with a concomitant enrichment of fatty acid metabolism and oxidation pathways in ACB1801-treated CT26 cells (**Figure 2C**). These results suggest that ACB1801 suppresses tumor cell glycolysis and support the hypothesis that it predisposes tumor cells to ferroptotic cell death by enhancing fatty acid metabolism and oxidation, thereby priming them for lipid peroxidation under oxidative stress conditions.

**Figure 2.**
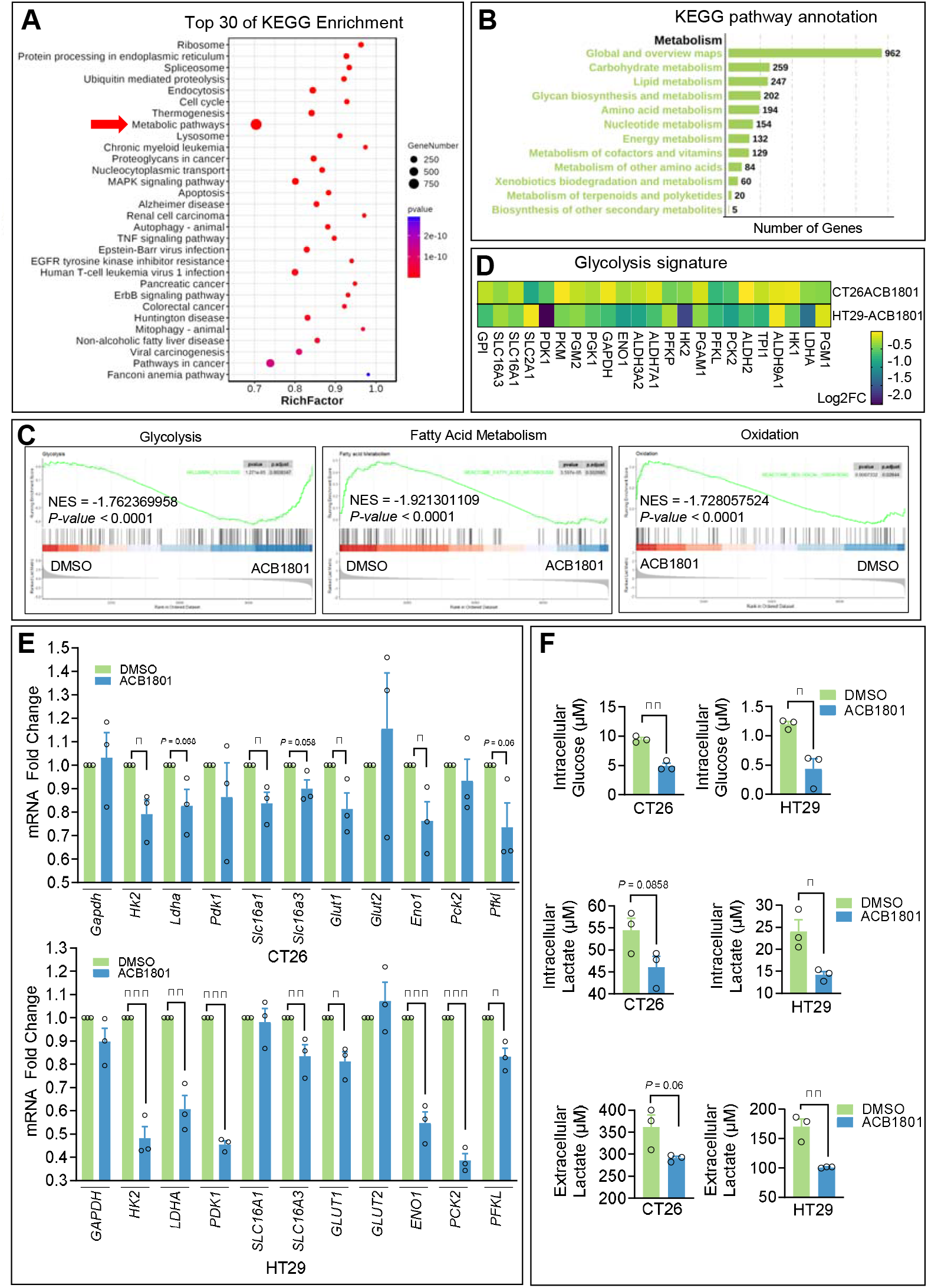
ACB1801 inhibits glycolytic and metabolic pathways in tumor and cancer cells. **(A)** KEGG pathway enrichment analysis of CD45-cells isolated from tumors treated with anti-PD-1 alone or with combination anti-PD-1/ACB1801. The dot plot shows the top 30 enriched KEGG pathways ranked by RichFactor, with dot size indicating the number of genes per pathway and color representing the adjusted p-value for significance (blue to red scale). Pathways associated with ACB1801-related functional changes in tumor CD45-cells are highlighted, including those related to metabolic processes. **(B)** KEGG pathway annotation of metabolic processes in CD45-cells from tumors treated with anti-PD-1 alone or in combination with ACB1801. The bar graph illustrates the number of genes involved in key metabolic pathways, grouped by category. The greatest gene representation is found in global and overview maps, with additional subdivisions including carbohydrate, lipid, glycan, amino acid, nucleotide, and energy metabolism, as well as biosynthetic and biodegradation pathways. This profile highlights metabolic processes most affected by ACB1801 treatment in the combination setting. **(C)** Gene set enrichment analysis (GSEA) comparing metabolic pathway activity between ACB1801- and DMSO-treated CT26 cells. The enrichment plots show normalized enrichment scores (NES) and p-values for glycolysis, fatty acid metabolism, and oxidation pathways. In all cases, metabolic pathway gene sets are significantly downregulated in ACB1801-treated cells relative to the DMSO control (p < 0.0001), indicating inhibition of glycolytic and oxidative metabolic programs by ACB1801. **(D)** Common glycolysis signatures in ACB1801-treated CT26 and HT29 cells. The heatmap displays log2 fold change (Log2FC) for key glycolytic genes in CT26 and HT29 cells following ACB1801 treatment. Each column represents a different gene implicated in glycolysis, while rows correspond to the 2 cell lines. Color scale indicates the degree of gene expression change, highlighting consistent downregulation of glycolytic pathways across both cell types in response to ACB1801. **(E)** ACB1801 suppresses glycolytic gene expression in CT26 and HT29 cells. The bar graphs show mRNA fold changes of key glycolytic genes in CT26 (top) and HT29 (bottom) cells treated with DMSO or ACB1801 (10 μM for CT26; 20 μM for HT29, 24 hours), as measured by quantitative RT-PCR. ACB1801 consistently downregulates multiple glycolytic genes, including HK2, LDHA, SLC16A1, PDK1, GLUT1, ENO1, and PFKL, in both cell lines. Data are presented as the fold change (FC) relative to the control (DMSO), with mean ± SEM from 3 independent experiments. Statistical significance was determined by an unpaired t-test (*p < 0.05, **p < 0.01, ***p < 0.001). **(F)** ACB1801 reduces intracellular glucose and lactate levels, as well as extracellular lactate secretion, in CT26 and HT29 cells. The bar graphs show the quantification of intracellular glucose, intracellular lactate, and extracellular lactate in cells treated with DMSO or ACB1801 (10 μM for CT26; 20 μM for HT29, 24hours). Both CT26 and HT29 cells exhibit significant decreases in intracellular glucose and lactate, along with reduced extracellular lactate release following ACB1801 treatment. Data represent the mean ± SEM from 3 independent experiments. Statistical significance was determined by an unpaired t-test (*p < 0.05; **p < 0.01).

We further explored the potential role of ACB1801 in suppressing tumor cell glycolysis and sensitizing tumor cells to ferroptosis. A comparative analysis identified 23 glycolytic genes commonly downregulated by ACB1801 in both mouse CT26 and human HT29 CRC cells, representing core enzymes involved in the 10 steps of the glycolytic pathway (**Figure 2D**). qRT-PCR validation in CT26 and HT29 cells further confirmed the downregulation of key glycolysis-related genes, including HK2, LDHA, SLC16A3, GLUT1, ENO1, and PFKL, following ACB1801 treatment (**Figures 2E**).

We next assessed the functional consequences of ACB1801-mediated downregulation of key glycolysis-related genes. Treatment with ACB1801 significantly reduced intracellular glucose and lactate levels as well as extracellular lactate in both CT26 and HT29 cells (**Figure 2F**), consistent with the coordinated downregulation of multiple glycolytic enzymes and transporters, including HK2, PFKL, ENO1, LDHA, and SLC16A3.

Similarly, when we assessed the expression of ferroptosis-related genes in mouse CT26 and human HT29 cells treated with ACB1801, we observed downregulation of several key ferroptosis suppressors, including SLC7A11, HSPA5, ATF4, NRF2, ENO1, and LCN2, along with upregulation of ferroptosis drivers such as ZEB1, STA1, and FADS2 (**Figure 3A**). qRT-PCR validation in CT26 and HT29 cells further confirmed the downregulation of SLC7A11, ATF4, NRF2, ENO1, LCN2 and HSPA5 following ACB1801 treatment (**Figure 3B**). These coordinated changes support the hypothesis that ACB1801 enhances the sensitivity of MSS CRC cells to ferroptotic cell death. To assess the functional consequences of the differentially regulated ferroptosis-related genes on ferroptosis sensitivity, we measured the GSH/GSSG ratio, a key indicator of cellular redox status [24]. Our results (**Figure 3C**) show a decrease in the GSH/GSSG ratio in ACB1801-treated CT26 and HT29 cells, indicating a shift toward increased cellular oxidative stress. We also observed a dose-dependent increase in reactive oxygen species (ROS) and lipid peroxidation in these cells following ACB1801 treatment (**Figure 3D**). Taken together, these findings strongly suggest that ACB1801 makes tumor cells vulnerable to ferroptotic cell death, a conclusion further supported by the observation that co-treatment with the specific ferroptosis inhibitor ferrostatin-1 completely abrogated ACB1801-dependent induction of cell death (**Figure 3E**).

**Figure 3.**
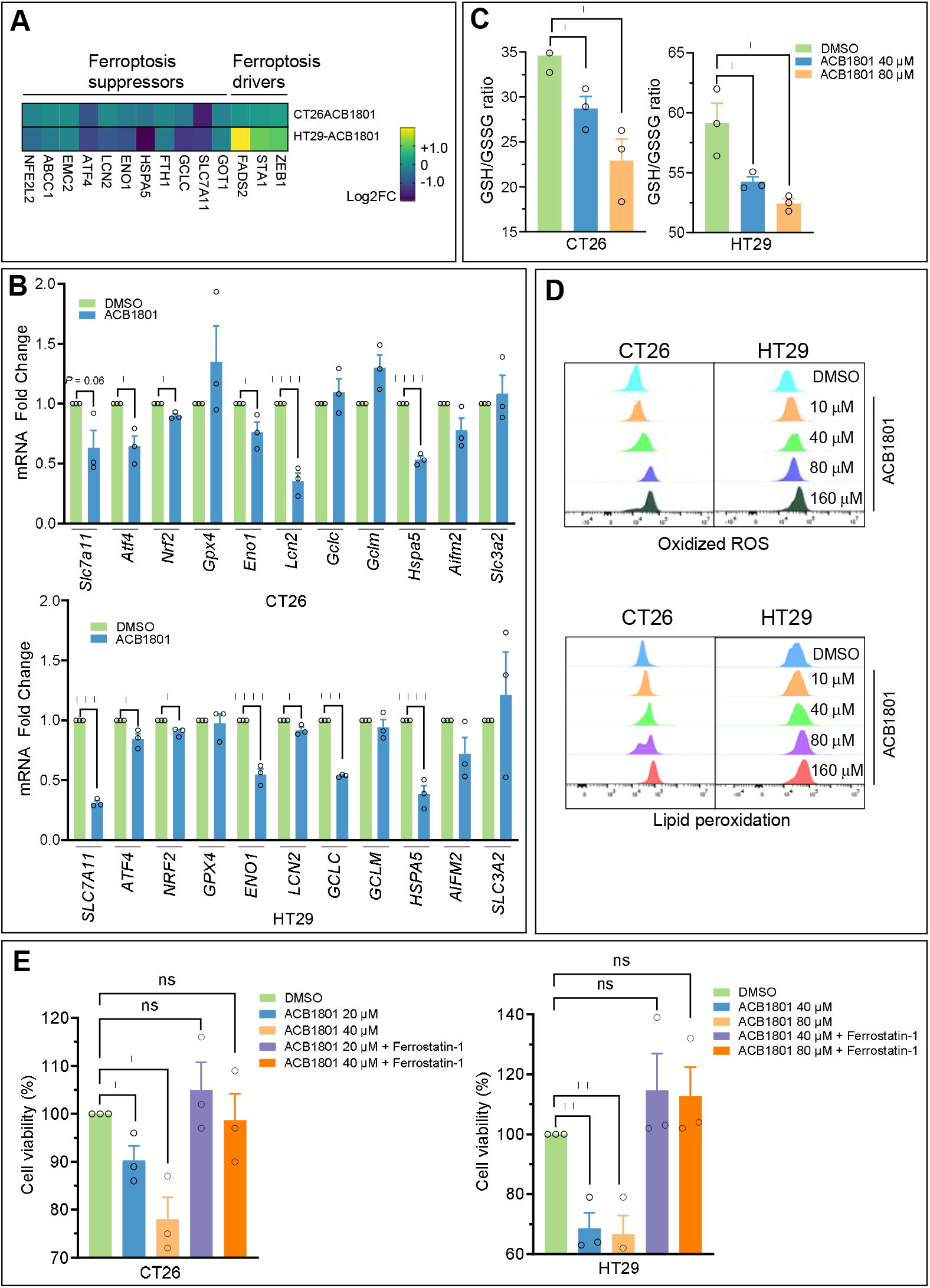
ACB1801 facilitates MSS CRC cells’ sensitivity to ferroptosis to modulate the tumor microenvironment. **(A)** Expression analysis of ferroptosis suppressor and driver genes in CT26 and HT29 cells following ACB1801 treatment (10 μM for CT26; 20 μM for HT29, 24hours). The heatmap illustrates log2 fold change (Log2FC) in the expression of key genes associated with ferroptosis suppression and induction for CT26 and HT29 cells. The color scale reflects relative up- or downregulation of each gene. Distinct expression patterns highlight the impact of ACB1801 on ferroptosis regulatory networks in both cell lines. **(B)** ACB1801 downregulates the expression of ferroptosis-related genes in HT29 cells. The bar graphs show mRNA fold changes of key genes involved in ferroptosis regulation following treatment with DMSO or ACB1801 (10 μM for CT26; 20 μM for HT29, 24hours), as measured by quantitative PCR. Significant decreases are observed for several ferroptosis suppressors and antioxidants, including SLC7A11, ATF4, NRF2, GPX4, ENO1, LCN2, GCLC, and GCLM. Data represent the mean ± SEM of the fold change relative to the DMSO-treated controls. Statistical significance was determined by an unpaired t-test (*p < 0.05; **p < 0.01; ***p < 0.001; ****p < 0.0001). **(C)** ACB1801 lowers the GSH/GSSG ratio in CT26 and HT29 cells. The bar graphs show the ratio of reduced glutathione (GSH) to oxidized glutathione (GSSG) in cells treated with DMSO or ACB1801 (40 μM and 80 μM). ACB1801 significantly decreases the GSH/GSSG ratio in a dose-dependent manner, indicating increased oxidative stress and reduced cellular antioxidant capacity. Data represent the mean ± SEM from 3 independent experiments. Statistical significance was determined by an unpaired t-test (*p < 0.05). **(D)** ACB1801 increases oxidized ROS and lipid peroxidation in CT26 and HT29 cells. The flow cytometry histograms show dose-dependent changes in oxidized reactive oxygen species (ROS, top) and lipid peroxidation (bottom) in CT26 and HT29 cells treated with escalating concentrations of ACB1801 compared to DMSO controls. Both cell lines exhibit increased oxidative stress and membrane lipid damage upon 24hours with ACB1801. **(E)** ACB1801 reduces cell viability via ferroptosis in CT26 and HT29 cells. The bar graphs show cell viability following treatment with DMSO or ACB1801 (20, 40, or 80 μM) in the absence or presence of the ferroptosis inhibitor ferrostatin-1. ACB1801 significantly decreases cell viability in both cell lines, an effect that is reversed by ferrostatin-1 co-treatment, indicating ferroptosis involvement. Data represent the mean ± SEM from 3 independent experiments. Statistical significance was determined by an ordinary one-way ANOVA (*p < 0.05; **p < 0.01; ns = not significant).

### ACB1801 increases the expression and release of CXCL10 in CT26 tumor cells via a STAT1-dependent mechanism

Given the elevated IFNγ signature in CT26 tumors treated with the ACB1801 and anti–PD-1 combination (**Supplementary Figure 2B**), and to determine whether ACB1801 directly influences cytokine and chemokine release from tumor cells, beyond the enrichment observed in CD45^−^ immune cells (**Supplementary Figure 2A**), we next profiled the cytokines and chemokines secreted by tumor cells in response to ACB1801. These factors likely contribute to reshaping the immune landscape in vivo, as shown in **Figure 1E**. We comprehensively profiled 111 cytokines and chemokines simultaneously using supernatants from untreated or ACB1801-treated cells. Under conditions of ACB1801-induced Tap1 upregulation (which served as an internal control), several cytokines and chemokines were differentially regulated, with CXCL10 notably upregulated and LCN2 and CXCL1 downregulated in ACB1801-treated cells compared to controls, (**Figure 4A**). We focused on CXCL10 due to its well-established role in promoting the infiltration of CXCR3+ CD8+ effector T cells into tumors in colorectal cancer models [25]. ELISA testing confirmed the increased release of CXCL10 in the supernatants of both mouse CT26 and human HT29 CRC cells treated with ACB1801 (**Figure 4B**). Furthermore, ACB1801 treatment also induced a significant increase in CXCL10 mRNA expression in these cells (**Figure 4C**), indicating that ACB1801 enhances not only CXCL10 release but also its gene expression. Notably, this increased CXCL10 release was observed not only in vitro, but also in the tumor microenvironment of CT26 tumors treated with ACB1801 alone, and to an even greater extent in those treated with the combination therapy (**Figure 4D**). GSEA of cultured CT26 cells revealed significant enrichment of the IFNg response signature following ACB1801 treatment (**Figure 4E**), suggesting activation of the STAT1 signaling pathway. Consistently, ACB1801 treatment increased STAT1 phosphorylation at Tyr701, pSTAT1(Y701) in both CT26 and HT29 cells, confirming the activation of STAT1 signaling (**Figure 4F**). We next assessed whether the upregulated expression of CXCL10 mRNA is a result of ACB1801-induced STAT1 activation, which is well-documented to drive CXCL10 expression [26]. Our findings in **Figure 4G and Supplementary Figure 3A** demonstrate that targeting STAT1 in both mouse CT26 and human HT29 cells abolished the ACB1801-induced increase in CXCL10. These results indicate that ACB1801 promotes CXCL10 overexpression and release in CRC cells through a STAT1-dependent mechanism.

**Figure 4.**
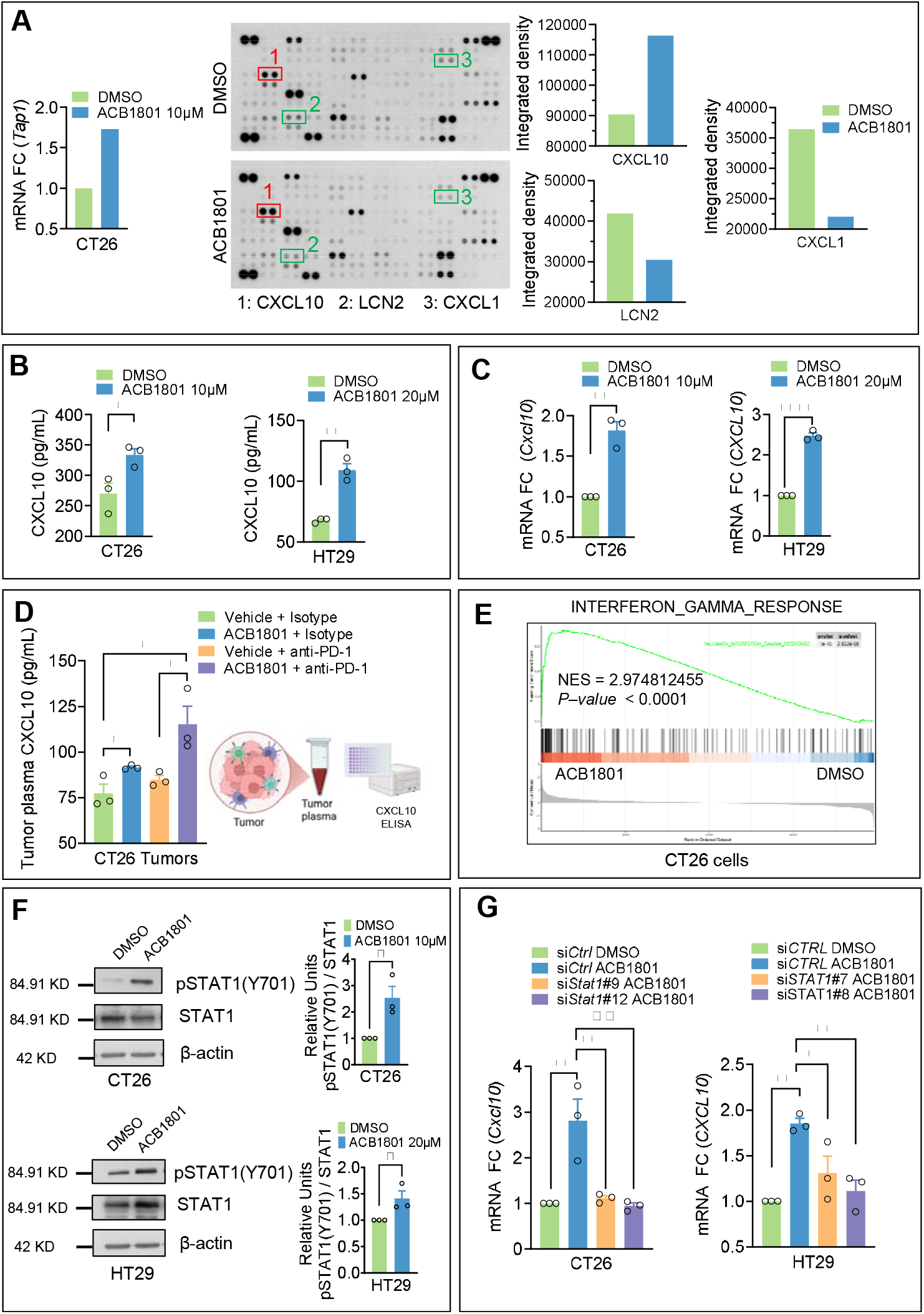
ACB1801 regulates CXCL10 expression in vitro and in vivo through STAT1. **(A)** Cytokine profiling of CT26 cell culture supernatants after treatment with DMSO or ACB1801 (10 μM) for 24 hours. The left panel shows an increase in Tap1 expression under these conditions. Each duplicate spot represents the relative abundance of an individual cytokine or chemokine. Highlighted boxes mark CXCL10 (1, red), LCN2 (2, green), and CXCL1 (3, green). The adjacent bar graphs show the integrated density quantification for CXCL10, LCN2, and CXCL1. ACB1801 treatment enhances CXCL10 secretion while reducing LCN2 and CXCL1 levels relative to the DMSO control. **(B and C)** Quantification of CXCL10 expression at the protein (B) and mRNA (C) levels following treatment with DMSO or ACB1801. (B) ELISA measurement of CXCL10 protein secretion in CT26 and HT29 cells treated with ACB1801 (10 μM for CT26; 20 μM for HT29) for 24 hours shows a significant increase in CXCL10 levels compared with DMSO. (C) Quantitative RT-PCR analysis of Cxcl10 (CT26) and CXCL10 (HT29) mRNA expression reveals elevated transcript levels in ACB1801-treated samples. Data represent the mean ± SEM from 3 independent experiments. Statistical significance was determined by an unpaired t-test (**p < 0.01, ***p < 0.001). **(D)** Quantification of CXCL10 levels in tumor plasma from CT26 tumors following the indicated treatments. CXCL10 concentrations were measured by ELISA. Data represent the mean ± SEM from 3 tumors per treatment group. Statistical significance was determined by an ordinary one-way ANOVA (*p < 0.05). **(E)** Gene set enrichment analysis (GSEA) of CT26 cells treated with ACB1801 versus DMSO control. ACB1801 treatment significantly enriches the interferon-gamma response gene signature, as reflected by a high normalized enrichment score (NES) and strong statistical significance (P < 0.0001). **(F)** Western blot analysis of STAT1 phosphorylation at Tyr701 (Y701) in CT26 and HT29 cells following 24 h treatment with DMSO or ACB1801 (10 µM for CT26; 20 µM for HT29). Whole-cell lysates prepared in RIPA buffer were immunoblotted for phospho-STAT1 (Y701), total STAT1, and β-actin as a loading control. Right panels show the ratio of phospho-STAT1 (Y701) to total STAT1, expressed as the mean ± SEM of three independent experiments. Statistical significance was assessed using an unpaired two-tailed t-test with Welch’s correction (p < 0.05). **(G)** Quantitative RT-PCR analysis of Cxcl10/CXCL10 mRNA expression in CT26 and HT29 cells after 24-hour treatment with DMSO or ACB1801 (10 μM for CT26; 20 μM for HT29), following transfection with control siRNA (siCTRL) or STAT1-targeting siRNAs (CT26: siStat11#9 and #12; HT29: siSTAT1#7 and #8). ACB1801-induced upregulation of Cxcl10/CXCL10 mRNA is attenuated upon STAT1 knockdown. Data are presented as the fold change (FC) relative to the control, with mean ± SEM from three independent experiments. Statistical significance was determined by an ordinary one-way ANOVA (*p < 0.05; **p < 0.01).

### ACB1801 enhances MHC-I expression and antigen presentation in MSS CRC through a STAT1/NLRC5-dependent mechanism

In addition to the metabolic pathways enriched in tumor cells isolated from combination-treated mice (ACB1801 + anti–PD-1) compared with anti–PD-1 monotherapy, KEGG pathway analysis (**Figure 2A**) also revealed significant enrichment of protein processing in the endoplasmic reticulum and ubiquitin-mediated proteolysis pathways. These pathways are closely linked to MHC-I antigen processing and presentation, supporting a potential role for ACB1801 in enhancing tumor antigen presentation. To further delineate this mechanism, we leveraged RNA-sequencing data to perform comparative transcriptomic profiling of untreated and ACB1801-treated CT26 cells, aiming to identify tumor cell–intrinsic processes regulating MHC-I antigen processing and presentation. Standardized data processing and differential gene expression (DEG) analysis (**Supplementary Figure 3B**) identified 116 significantly regulated genes, including 85 upregulated and 31 downregulated transcripts following ACB1801 treatment.

By applying PPI network construction on DEG, our data revealed that protein processing is the most regulated process upon ACB1801 treatment (**Figure 5A**). GSEA further revealed enriched antigen processing and presentation pathways in CT26 cells treated with ACB1801 (**Figure 5B)**. This finding is further supported by data showing that ACB1801 treatment upregulates the MHC-I genes *Tap1, Tap2*, and *B2M* in both CT26 mouse and HT29 human MSS CRC cells (**Supplementary Figure 4A**) consistent with our previous observations in melanoma cells [20]. As a result, ACB1801-treated mouse and human CRC cells exhibited increased surface expression of H-2K^d^ and HLA-A, B, C molecules, compared to untreated cells (**Figure 5C**). The functional significance of ACB1801-induced upregulation of MHC-I was demonstrated by OVA-derived SIINFEKL peptide loading on MHC-I in MC38 CRC cells (**Figure 5D**), which subsequently enhanced OT-I CD8^+^ T cell-mediated killing of OVA-loaded MC38 cells (**Figure 5E)**.

**Figure 5.**
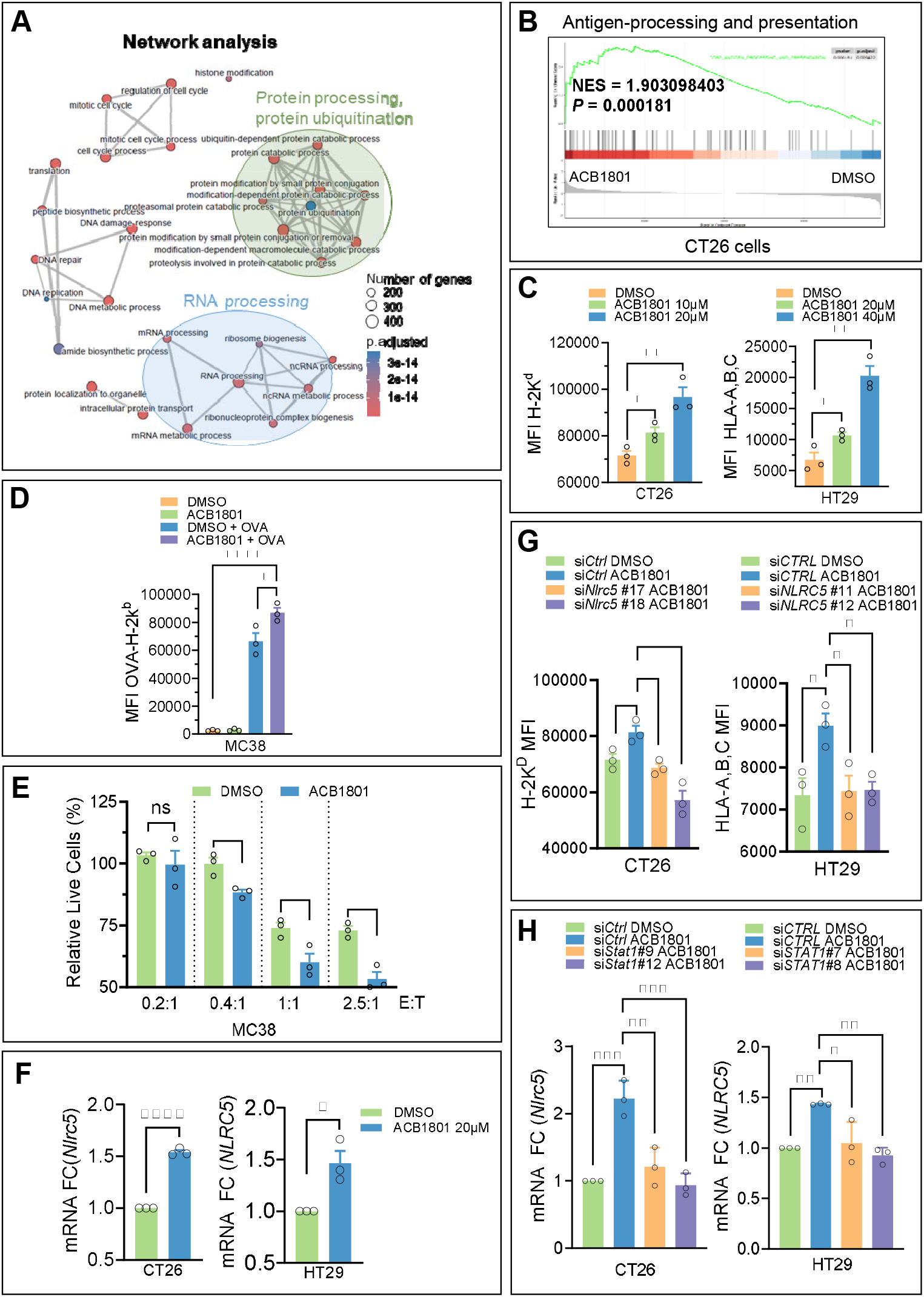
ACB1801 upregulates antigen processing and presentation pathways through STAT1-dependent induction of NLRC5. **(A)** Gene ontology (GO) enrichment network analysis of differentially expressed genes in CT26 tumors treated with ACB1801. Network nodes correspond to enriched GO biological processes, with node size proportional to the number of associated genes and edges indicating functional relationships between GO terms. Adjusted p-values are color-coded to reflect enrichment strength. Major functional clusters center on protein modification, ubiquitin-dependent protein catabolism, and protein processing (highlighted in blue), consistent with coordinated regulation of pathways involved in antigen processing and presentation. **(B)** Gene set enrichment analysis (GSEA) comparing CT26 tumors treated with ACB1801 versus DMSO reveals significant enrichment of the antigen processing and presentation pathway. The enrichment plot shows a high normalized enrichment score (NES = 1.90) and strong statistical significance (p = 0.000181), indicating robust upregulation of antigen-processing genes following ACB1801 treatment. **(C)** ACB1801 increases cell-surface MHC-I expression in CT26 and HT29 cells. The representative flow cytometry histograms show H-2Kd (CT26) and HLA-A, B, and C (HT29) expression after treatment with DMSO or ACB1801 (20 μM, 40 μM) for 24 hours. The bar graphs represent the mean fluorescence intensity (MFI) from 3 independent experiments ± SEM. Statistical significance was determined by an unpaired t-test: *p < 0.05, **p < 0.01. **(D)** ACB1801 enhances the presentation of the OVA_257–264 (SIINFEKL) antigen in MC38 cells. Cells were pulsed with SIINFEKL peptide for 8 hours, followed by treatment with either DMSO or ACB1801 for 24 hours. Surface OVA–H-2Kb complexes were quantified by flow cytometry using PE–OVA–H-2Kb staining. The representative flow cytometry histograms (top) and bar graphs (bottom) show the mean fluorescence intensity (MFI) from 3 independent experiments ± SEM. Statistical significance was determined by an unpaired t-test (*p < 0.05; ****p < 0.0001), demonstrating that ACB1801 significantly increases SIINFEKL presentation. **(E)** ACB1801 enhances OT-I CD8+ T cell-mediated killing of MC38-OVA tumor cells. MC38-OVA cells were pulsed with SIINFEKL peptide for 8 hours and co-cultured with activated OT-I CD8+ T cells at the indicated effector: target (E:T) ratios in the presence of DMSO or ACB1801. After 48 hours, tumor cell numbers were quantified using Precision Count Beads. The bar graphs show MC38-OVA cell viability (% ± SEM) from 3 independent experiments. Statistical significance was determined by unpaired t-test (ns = not significant; *p < 0.05; ***p < 0.001), demonstrating that ACB1801 significantly enhances OT-I-mediated cytotoxicity, particularly at higher E:T ratios. **(F)** Quantitative RT-PCR analysis of NLRC5 mRNA levels in CT26 and HT29 cells after treatment with ACB1801 (20 μM) or DMSO for 24 hours. Data are presented as mean fold change (FC) relative to DMSO-treated samples from 3 independent experiments (± SEM). Statistical significance was determined by an unpaired t-test (*p < 0.05; ****p < 0.0001). **(G)** Flow cytometry analysis of cell surface MHC-I expression in CT26 and HT29 cells following 24-hour treatment with ACB1801 (20 μM) or DMSO. Cells were transfected with either control siRNA (siCtrl) or NLRC5-targeted siRNAs (siNLRC5#17 and #18 for CT26; siNLRC5#11 and #12 for HT29). Representative flow cytometry histograms show H-2KD (CT26, APC conjugated) and HLA-A, B, C (HT29, PE conjugated) staining. The bar graphs show mean fluorescence intensity (MFI), presented as mean ± SEM from 3 independent experiments. Statistical significance was calculated using an ordinary one-way ANOVA : **p < 0.01, *p < 0.05. **(H)** Quantitative RT–PCR analysis of NLRC5 expression in CT26 and HT29 cells following STAT1 knockdown and ACB1801 treatment. Cells were transfected with either control siRNA (siCtrl) or STAT1-targeting siRNAs (CT26: siSTAT1#9 and #12; HT29: siSTAT1#7 and #8), then treated for 24 hours with DMSO or ACB1801. NLRC5 mRNA levels are presented as the fold change (FC) relative to the control. Data represent the mean ± SEM from 3 independent experiments. Statistical significance was determined by an ordinary one-way ANOVA (*p < 0.05, **p < 0.01, ***p < 0.001).

MHC-I antigen processing and presentation are regulated by various pathways and transcription factors, with NLRC5 protein identified as a key transcriptional regulator of MHC-I-related genes [27]. Our data in **Figure 5F** demonstrates that ACB1801 significantly increases NLRC5 expression in both CT26 mouse and HT29 human CRC cells. The ACB1801-induced increase in the expression of H-2K⍰ and HLA-A, -B, and -C, as well as the MHC-I-related genes TAP1 and TAP2, requires NLRC5, as this increase was not observed in cells transfected with NLRC5 siRNA (**Figure 5G and Supplementary Figure 4B**). These findings further demonstrate that ACB1801 enhances MHC-I through an NLRC5-dependent mechanism in MSS CRC cells.

NLRC5 expression is regulated by various inflammatory signaling pathways, including type I and type II interferons (IFNs) through the activation of STATs [28]. We therefore postulated that the ACB1801-dependent increase in NLRC5 may involve an activation of the IFNγ pathway via STAT signaling. This theory is supported by our RNA-seq data derived from untreated and ACB1801-treated CT26 cells, which revealed significant enrichment of the IFNγ response and the increased pSTAT1(Y701) phosphorylation level in treated cells (**Figure 4E and 4F**). Consistent with these findings, **Figure 5H** shows that treatment of CT26 mouse and HT29 human CRC cells with ACB1801 increases NLRC5 mRNA levels, and targeting STAT1 abolishes the ACB1801-dependent increase in NLRC5 (**Supplementary Figure 4B and Figure 3E**). Together, these findings provide compelling evidence that ACB1801 triggers an early event involving STAT1 phosphorylation, positioning pSTAT1 as a key upstream regulator of NLRC5 expression and MHC-I antigen processing and presentation.

## Discussion

In this study, we demonstrate that the harmine derivative ACB1801 synergizes with and enhances the efficacy of anti-PD-1 therapy in the MSS CRC CT26 model, extending our previous observations in melanoma [20]. More importantly, we provide mechanistic insights into how ACB1801 primes tumor cells and the tumor microenvironment to improve responsiveness to anti-PD-1 therapy. Notably, ACB1801 enhances IFNγ responsiveness and upregulates MHC-I expression, thereby priming tumor cells for immune recognition. However, these effects alone are insufficient to trigger a robust anti-tumor response, as treatment with ACB1801 alone did not significantly inhibit tumor growth, likely because T cell activity remains suppressed by immune checkpoint pathways. When ACB1801 is combined with anti-PD-1, which overcomes this inhibition, the primed tumor microenvironment translates into a robust immune response characterized by an increased frequency of CD8+ T cells and a reduced frequency of regulatory T cells, resulting in a significantly higher CD8/Treg ratio within the TME.

Our transcriptomic analysis indicates that the priming effect of ACB1801 on tumor cells is primarily driven by enhanced IFNγ responsiveness, characterized by early STAT1 activation that subsequently engages the NLRC5/CXCL10 axis. Specifically, we observed that ACB1801 treatment upregulates a cluster of interferon-stimulated genes (ISGs), including IRF7, STAT2, DDX58, and OAS1/2, which collectively amplify the IFNγ response and reinforce downstream immune signaling. Several plausible, non-mutually exclusive mechanisms may underlie ACB1801’s ability to activate STAT1 in tumor cells. ACB1801 downregulates key genes known to repress IFNγ signaling, thereby promoting STAT1 activation. Our transcriptomic analysis of ACB1801-treated CT26 cells revealed a significant downregulation of TRIB3 (**Supplementary Figure 3B**), a gene well described as repressing the STAT1–CXCL10 axis, reducing CD8^+^ T-cell infiltration, and promoting immune evasion in colorectal cancer [29]. Whether this effect of ACB1801 is linked to its properties as a β-carboline alkaloid derived of harmine [10], as known inhibitor of DYRK1A [30], remains to be determined.

Another potential mechanism is that ACB1801 promotes the autocrine secretion of chemokines, such as CXCL10, thereby establishing a positive feedback loop that sustains STAT1 activation. This mechanism is supported by our transcriptomic data, which show that ACB1801 downregulates DUSP5, a nuclear phosphatase that specifically dephosphorylates MAPK (ERK1/2) [31]. Downregulation of DUSP5 leads to increased ERK1/2 phosphorylation, which in turn promotes CXCL10 expression through transcription factors such as AP-1 and NF-κB [32]. This upregulation of CXCL10 reinforces the autocrine loop, further sustaining STAT1 activation. Collectively, ACB1801 appears to engage multiple overlapping mechanisms that converge on enhanced IFN responsiveness, STAT1 activation, NLRC5 upregulation, and increased MHC-I expression.

One important consequence of STAT1 activation by ACB1801 is the modulation of the chemokine network released by tumor cells, which shapes the tumor immune microenvironment. The simultaneous upregulation of CXCL10 and downregulation of LCN2 upon ACB1801 treatment may represent a synergistic mechanism driving the remodeling of the immune landscape and regulating tumor cell ferroptosis. We believe that CXCL10 upregulation is likely established a chemotactic gradient promoting CD8+ T-cell infiltration [8, 33], while reduced LCN2 expression, as shown in **Supplementary Figure 5A**, enhanced tumor cell susceptibility to ferroptosis, as previously reported in colon cancer cells [34]. Although the molecular mechanisms underlying CXCL10 upregulation via the STAT1/NLRC5 axis are well established [35], the mechanisms responsible for LCN2 downregulation in this context remain to be elucidated.

The reduction in Treg frequency in tumors treated with ACB1801 and anti-PD-1 combination therapy may be mechanistically linked to ACB1801-mediated downregulation of LCN2 as demonstrated in **Supplementary Figure 5**. Recent work has shown that tumor-derived LCN2 promotes intratumoral Treg accumulation by binding SLC22A17 on macrophages, driving their polarization toward an immunosuppressive phenotype that supports Treg recruitment and stability [36]. Although ACB1801 downregulates LCN2, a significant reduction in Treg frequency was observed only with combination therapy, not with ACB1801 alone. This suggests that LCN2 suppression primarily limits de novo Treg recruitment, whereas established Tregs, characterized by high PD-1 expression, require PD-1 blockade for their destabilization and functional impairment. Thus, restricting Treg influx via LCN2 suppression while disrupting Treg maintenance through anti–PD-1 likely explains the synergistic increase in the CD8^+^/Treg ratio observed in the combination group.

Another important and unexpected function of ACB1801 is its ability to reprogram tumor cell metabolism. This metabolic shift is characterized by decreased intracellular glucose and reduced extracellular lactate levels, driven by the coordinated downregulation of key glycolytic enzymes and transporters [37]. Such reprogramming has significant therapeutic relevance in colorectal cancer, where MSS CRC tumors are highly dependent on aerobic glycolysis. The resulting accumulation of lactate acidifies the TME and profoundly impairs CD8+ T-cell proliferation, cytokine secretion, and cytotoxic activity [38, 39], while concurrently promoting Treg differentiation and stability [40]. By lowering intracellular glucose, reducing lactate production, and decreasing the expression of key ferroptosis suppressor genes, we believe that ACB1801 alleviates these immunosuppressive pressures, restores CD8+ T-cell fitness, and increases tumor cell vulnerability to ferroptosis; these factors collectively shift the TME toward a more immune-permissive state.

Several lines of evidence from our study support this mechanism. First, transcriptomic profiling of CD45+ cells isolated from tumors treated with ACB1801 + anti-PD-1 revealed enrichment of adaptive immune activation, cytotoxic pathways, and IFNγ signaling, consistent with a more metabolically favorable niche for effector T cells. Second, analyses of TCGA CRC datasets showed that ferroptosis suppressor genes and glycolysis signature genes were negatively correlated with CD8+ T-cell markers and positively correlated with Treg markers (**Supplementary Figure 5B**), supporting the metabolic and immunological remodeling effect of ACB1801. Ferroptosis can elicit features of immunogenic cell death, as lipid peroxidation–driven membrane damage promotes the release of danger-associated molecular patterns (DAMPs) and inflammatory mediators that can enhance dendritic cell activation and antitumor immune responses [41]. Together, these findings support a dual mechanism for ACB1801: enhancing tumor immunogenicity through increased IFN responsiveness and MHC-I expression, while simultaneously reprogramming tumor metabolism to counteract glycolysis- and lactate-driven immunosuppression (**Figure 6**). By integrating these effects, ACB1801 promotes a more immune-supportive TME and increases tumor susceptibility to cytotoxic T-cell attack.

**Figure 6:**
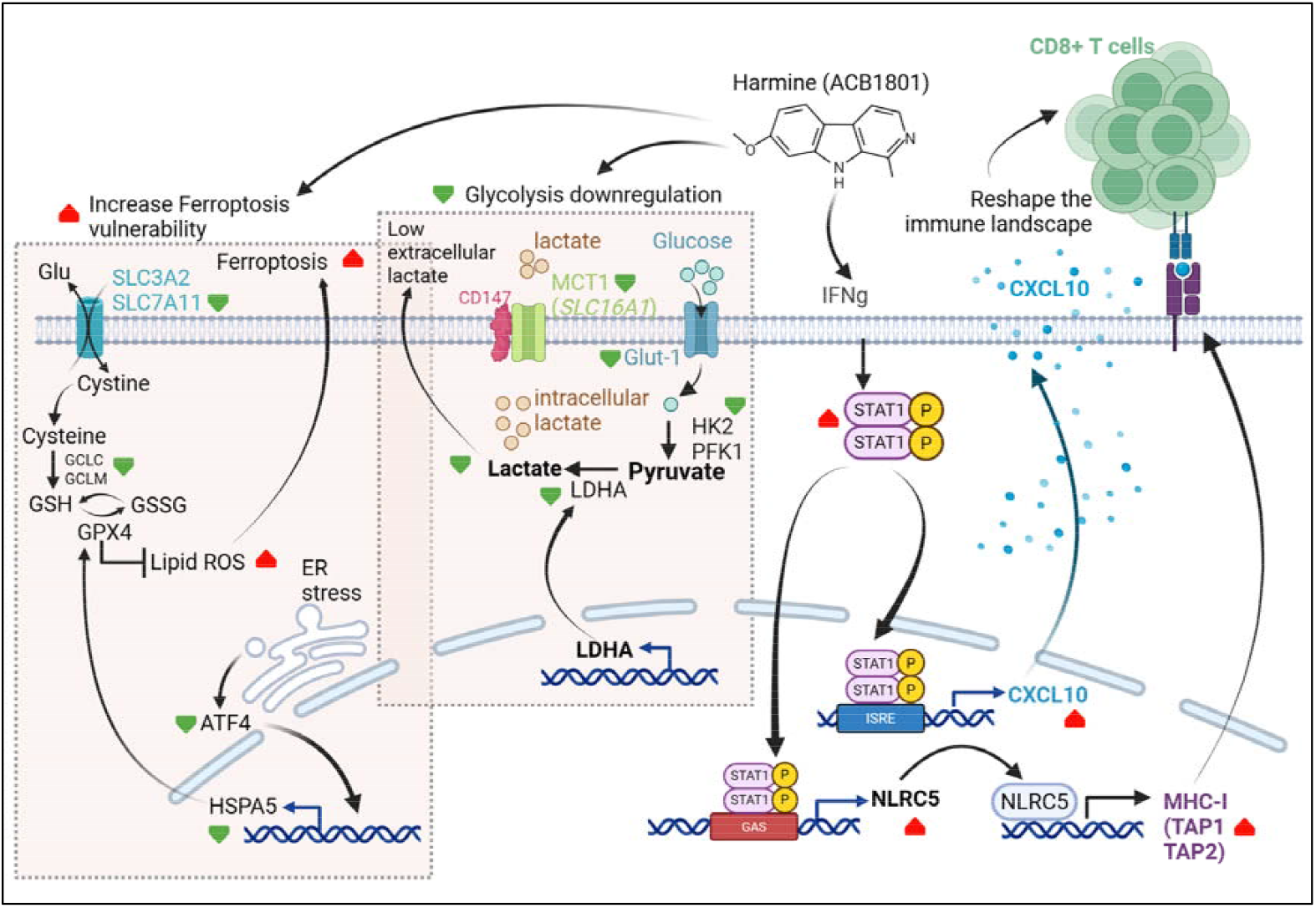
Proposed mechanism of action of ACB1801. ACB1801 enhances tumor immunogenicity by activating the IFNg /STAT1 signaling pathway, leading to the upregulation of NLRC5 and CXCL10. Increased NLRC5 expression promotes the transcription of MHC class I–related genes (including TAP1 and TAP2), resulting in enhanced MHC-I surface expression and improved antigen loading. In parallel, STAT1 activation induces CXCL10 expression and secretion, thereby reshaping the tumor immune microenvironment through the recruitment of CD8+ T cells. ACB1801 also reprograms tumor cell metabolism to counteract glycolysis- and lactate-driven immunosuppression. This effect is associated with the downregulation of key glycolytic and lactate transport genes, including SLC16A1 (MCT1), GLUT1, HK2, PFK1, and LDHA, leading to reduced intracellular and extracellular lactate levels. In addition, ACB1801 increases tumor cell vulnerability to ferroptosis by downregulating several genes directly or indirectly involved in redox homeostasis and ferroptotic resistance, including SLC7A11, GCLC/GCLM, ATF4, and HSPA5. Suppression of the SLC7A11–GCLC–GPX4 axis impairs glutathione-dependent detoxification of lipid peroxides, resulting in enhanced lipid ROS accumulation. Furthermore, inhibition of the ATF4/HSPA5 stress-response pathway further alleviates GPX4-mediated protection, collectively amplifying ferroptotic susceptibility.

## Conclusions

This study demonstrates that the harmine derivative ACB1801 overcomes anti-PD-1 resistance in MSS colorectal cancer through a dual mechanism: enhancing tumor immunogenicity via STAT1-dependent activation of the NLRC5/CXCL10 axis, and inducing metabolic vulnerability by suppressing glycolysis and sensitizing tumor cells to ferroptosis. The combination of ACB1801 with anti-PD-1 significantly inhibited tumor growth and improved survival, accompanied by increased CD8+ T cell infiltration and reduced regulatory T cells in the tumor microenvironment. These findings address a critical unmet clinical need, as MSS CRC represents approximately 85% of colorectal cancer cases yet remains largely refractory to current immunotherapies. By converting immunologically “cold” tumors into “hot,” immunoresponsive tumors through complementary mechanisms, ACB1801 offers a promising therapeutic strategy for this substantial patient population. Our data provide a strong preclinical rationale for clinical evaluation of Harmine derivative in combination with immune checkpoint blockade for MSS CRC patients who currently lack effective immunotherapeutic options.

## List of Abbreviations

*ICB*: Immune checkpoint blockade
*TME*: Tumor microenvironment
*TILs*: Tumor-infiltrating lymphocytes
*PD-1*: Programmed cell death protein 1
*MSS*: Microsatellite stable
*CRC*: Colorectal cancer
*dMMR*: Deficient mismatch repair
*MSI-H*: High microsatellite instability
*CTLs*: Cytotoxic T lymphocytes
*TMB*: Tumor mutational burden
*MHC-I*: Major histocompatibility complex class I
*APCs*: Antigen-presenting cells
*CXCL10*: C-X-C motif chemokine ligand 10
*DYRK1A*: Dual-specificity tyrosine-(Y)-phosphorylation regulated kinase 1A
*Tregs*: Regulatory T cells
*DEG*: Differentially expressed genes
*PPI*: Protein–protein interaction
*GSEA*: Gene set enrichment analysis
*OVA*: Ovalbumin
*NLRC5*: NOD-like receptor family CARD domain containing 5
*COAD*: Colon adenocarcinoma
*ROS*: Reactive oxygen species
*GSH*: Glutathione
*GSSG*: Glutathione disulfide.

## Declarations

### Ethics approval and consent to participate

All animal experiments were conducted in accordance with European Union guidelines. The in vivo protocols were approved by the Ethical Committee of the Luxembourg Institute of Health, the Animal Welfare Society, and the Luxembourg Ministry of Agriculture, Viticulture and Rural Development (Approval No. LECR-2018-12).

### Consent for publication

Not applicable

### Availability of data and materials

The sequencing files are deposited on GEO database (https://www.ncbi.nlm.nih.gov/geo/browse/) with the accession numbers: GSE (To be included upon acceptance).

### Author Contributions

BJ, CA, AI, MM, ED, GB and RG initiated the project and designed the research plan. RG, KVM, EB and MGT performed the in vitro experiments. CP, AO, RG, MP, TLR, DM and KVM carried out the in vivo experiments under the supervision of BJ. RG and CL conducted the bioinformatics analysis. Data analysis was performed by BJ, RG, AI and CA. The manuscript was written by BJ and RG and revised with input from all authors.

## Acknowledgements

We thank AC-Bioscience for generously providing the ACB1801 compound. We also acknowledge the LIH Animal Core Facility for their valuable support with the animal experiments.

## Figures and Legends for supplementary figures

**Supplementary Figure 1.**
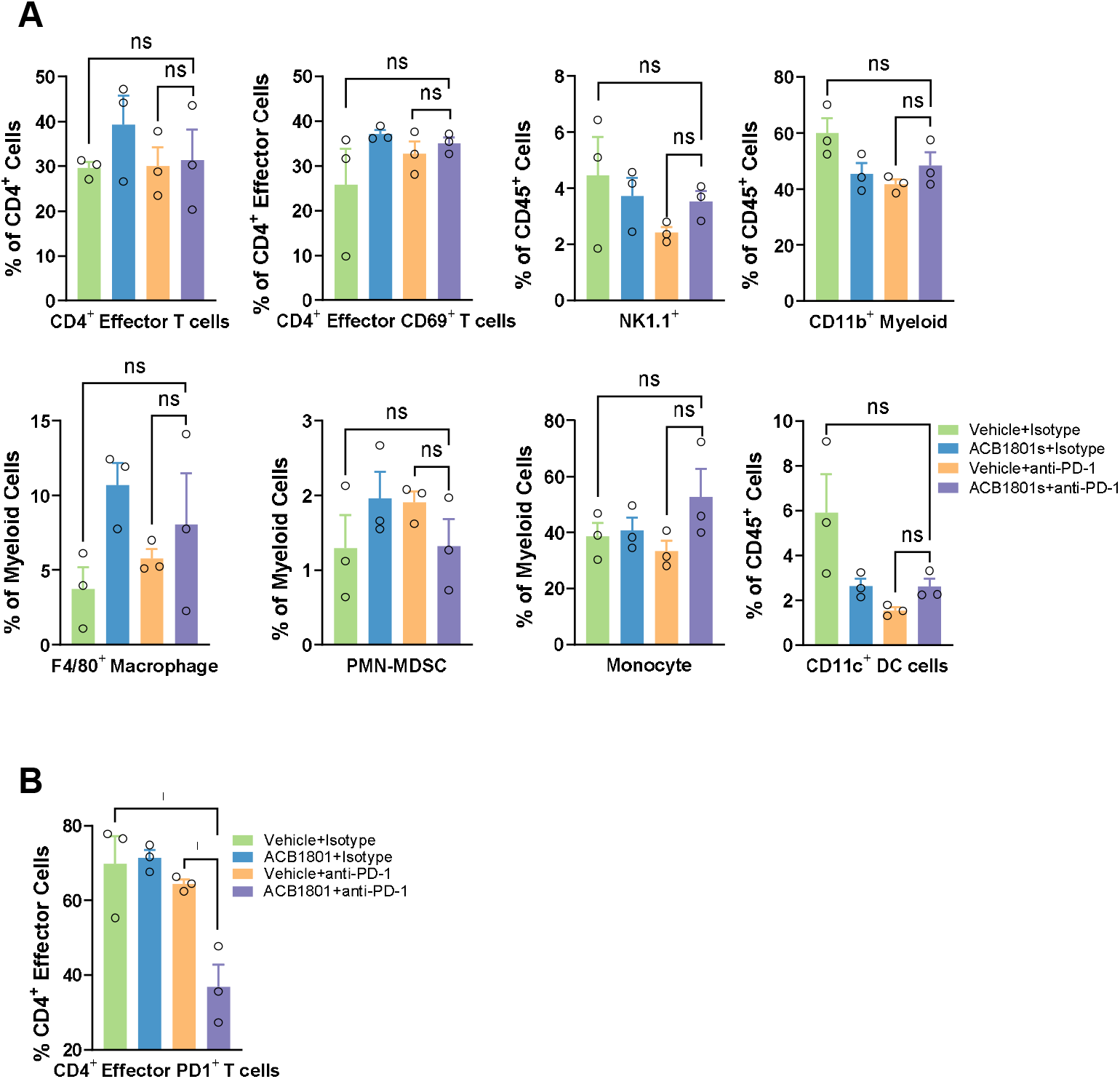
**(A)** Flow cytometric profiling of tumor-infiltrating immune cells in CT26 tumor-bearing mice treated as indicated. Quantified populations include CD4+ effector T cells, CD4+ CD69+ effector T cells, NK1.1+ cells, CD11b+ myeloid cells, F4/80+ macrophages, PMN-MDSCs, monocytes, and CD11c+ dendritic cells within CD45+ or myeloid compartments. Data represent the mean ± SEM of 3 tumors from each group. Statistical significance was determined by an ordinary one-way ANOVA (ns = not significant). **(B)** Percentage of CD4+ PD-1+ effector T cells among total CD4+ effector T cells across treatment groups. A significant reduction in CD4+ PD-1+ effector T cells was observed in the ACB1801+anti-PD-1 combination group compared to controls. Data represent the mean ± SEM of three tumors from each group. Statistical significance was determined by an ordinary one-way ANOVA (*p < 0.05).

**Supplementary Figure 2.**
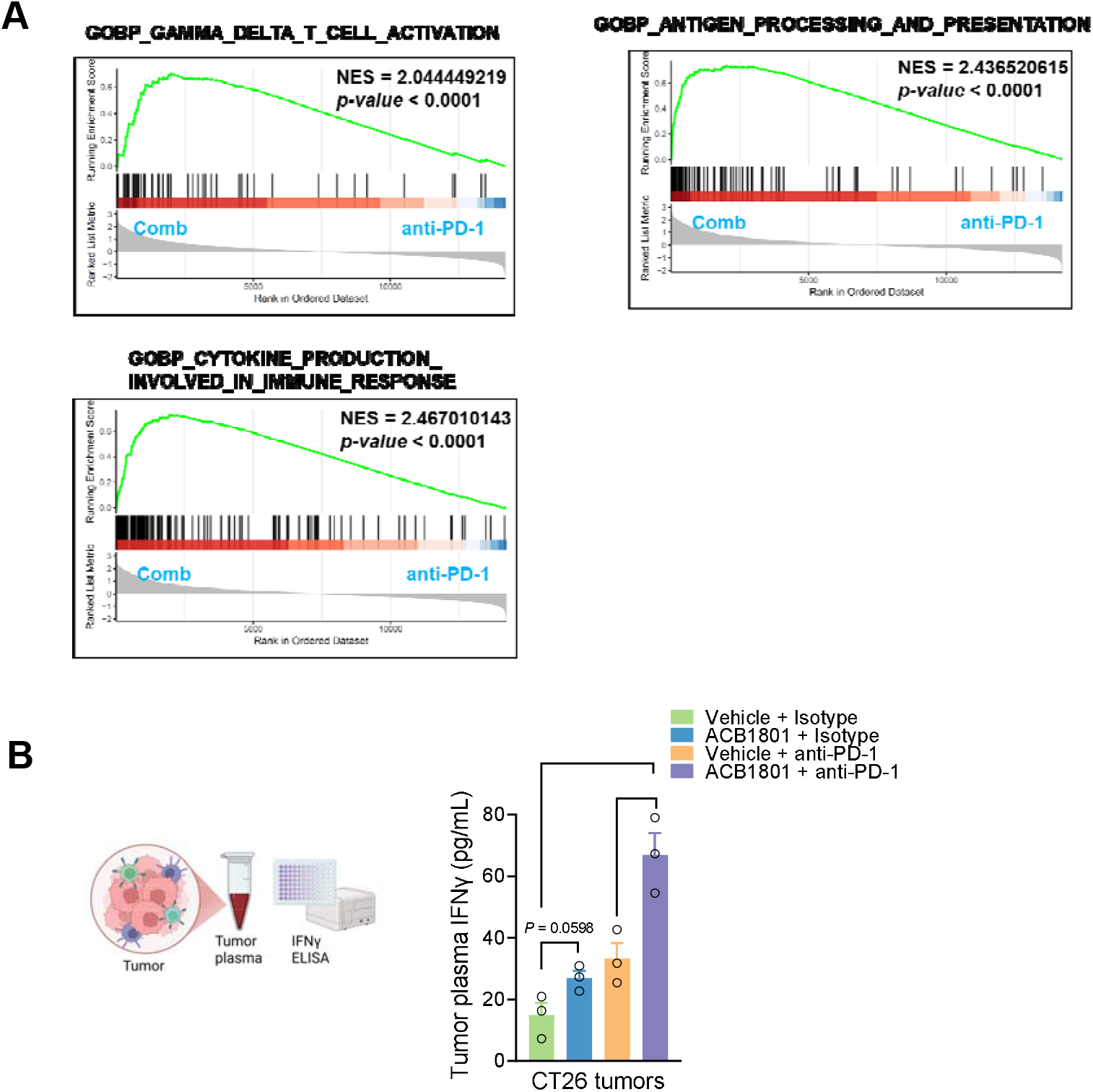
**(A)** Gene set enrichment analysis (GSEA) of RNA-seq data from isolated intratumoral CD45+ cells comparing the combination (combo) treatment group (ACB1801 + anti-PD-1) with the single-agent anti-PD-1 group. The enrichment plots show significant upregulation of gene sets associated with gamma/delta T cell activation, cytokine-production involved in immune response and antigen processing and presentation in the combination group. For each pathway, the green line represents the running enrichment score, vertical black bars indicate the positions of genes from the gene set in the ranked list, and the normalized enrichment score (NES) and p-value are reported within each panel. **(B)** Quantification of intratumoral IFNγ secretion in CT26 tumor-bearing mice. Tumors were enzymatically dissociated, and tumor plasma was isolated and concentrated by centrifugation. IFNγ levels were measured using a mouse IFNγ ELISA kit. The bar graphs represent the mean IFNγ concentration (pg/mL) ± SEM from 3 tumors per treatment group. Statistical significance was determined by an ordinary one-way ANOVA (*p < 0.05; **p < 0.01).

**Supplementary Figure 3.**
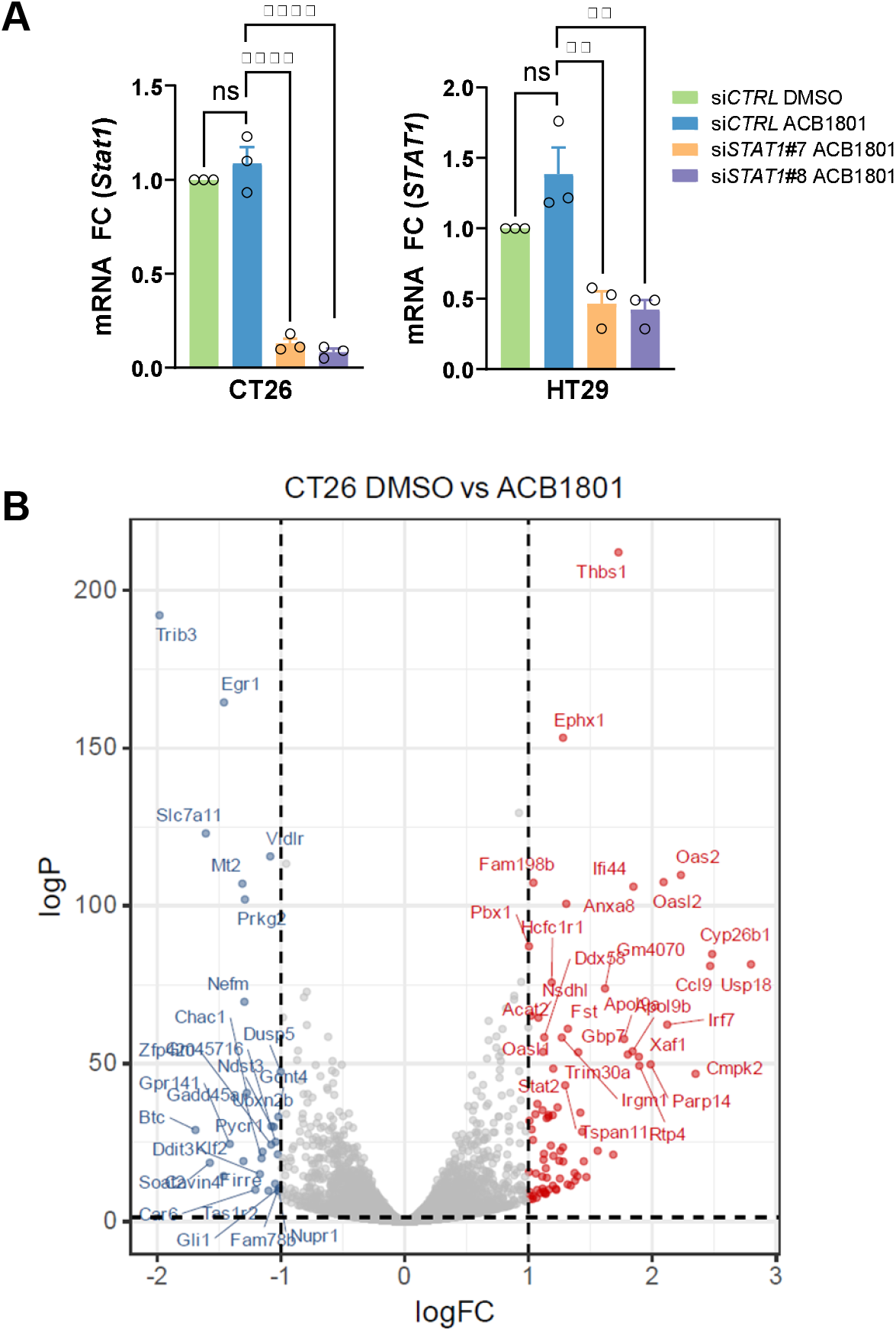
**(A)** Depletion of STAT1 with siRNAs targeting mouse *Stat1* or human *STAT1*. CT26 cells were transfected with siCTRL or 2 siRNAs targeting mouse Stat1 (#7 and #8), and HT29 cells were transfected with siCTRL or 2 siRNAs targeting human STAT1 (#7 and #8). Following treatment with DMSO or ACB1801, Stat1/STAT1 mRNA levels were quantified by RT-PCR. Data represent the mean ± SEM of the fold change (FC) relative to the siCTRL DMSO from 3 independent experiments. Statistical significance was determined by an ordinary one-way ANOVA (ns, not significant; **p < 0.01; ****p < 0.0001). **(B)** Volcano plot showing differentially expressed genes in CT26 tumors treated with ACB1801 compared to DMSO. Each point represents an individual gene, with log2 fold change (logFC) on the x-axis and −log10(p-value) on the y-axis. Upregulated genes in ACB1801-treated tumors are shown in red and downregulated genes in blue; no significant genes are shown in gray. Dashed vertical lines indicate log2FC cutoffs (±1), and the dashed horizontal line marks the p-value significance threshold.

**Supplementary Figure 4.**
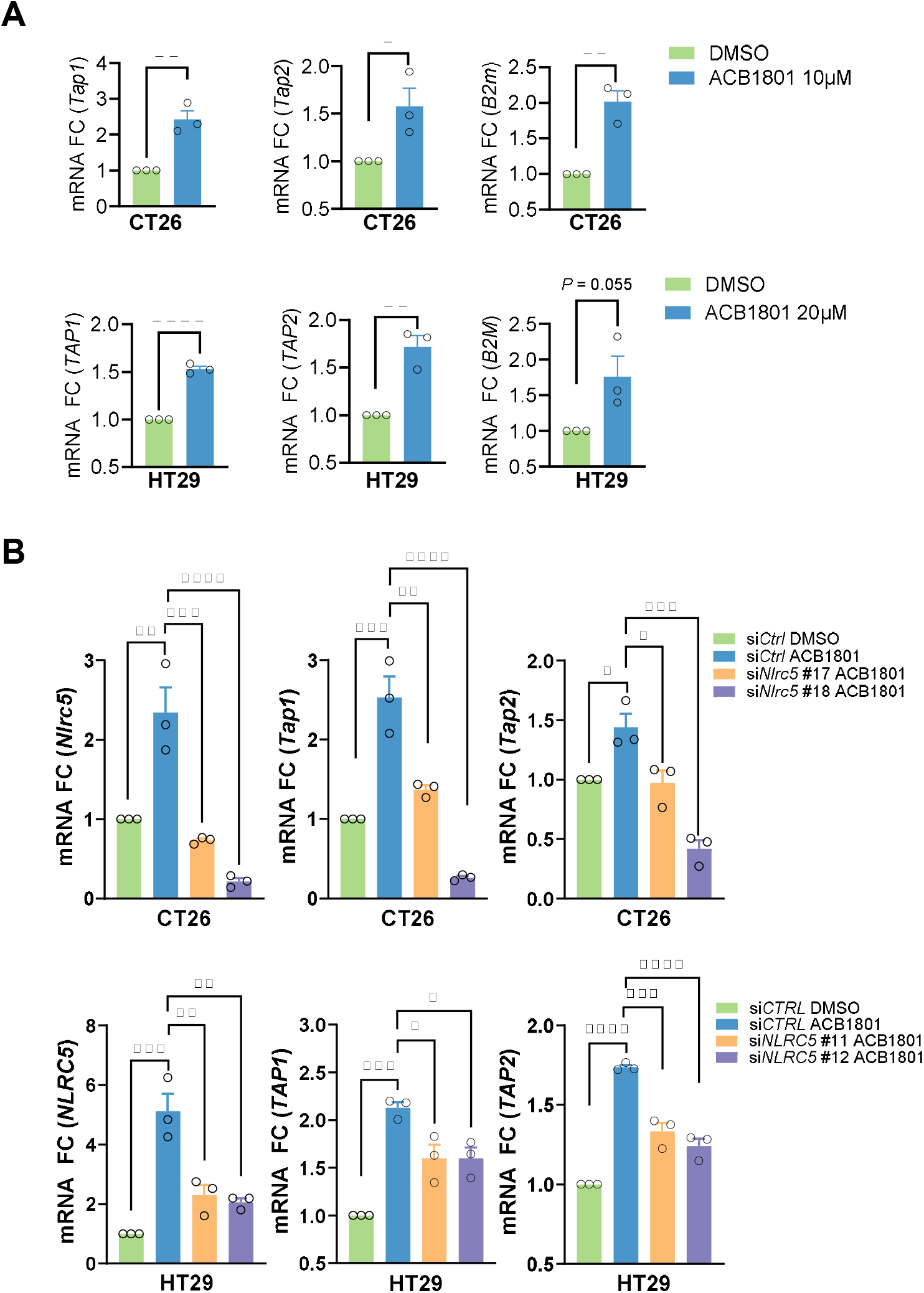
**(A)** ACB1801 enhances the expression of MHC-I–associated genes in murine (CT26) and human (HT29) cell lines. Cells were treated with ACB1801 (10 μM) or DMSO for 24 hours, and mRNA levels of Tap1/TAP1, Tap2/TAP2, and B2m/B2M were quantified by RT-PCR and normalized to the control. The bar graphs represent the mean fold change from 3 independent experiments ± SEM. Statistical significance was determined by an unpaired t-test: *p < 0.05, **p < 0.01, ****p < 0.0001. **(B)** Depletion of NLRC5 blocks the ACB1801-mediated induction of MHC-I–related genes. CT26 cells were transfected with control siRNA (siCTRL) or two siRNAs targeting mouse Nlrc5 (siNlrc5 #11 or #18), and HT29 cells were transfected with siCTRL or two siRNAs targeting human NLRC5 (siNLRC5 #11 or #12). Following transfection, cells were treated with DMSO or ACB1801, and mRNA levels of Nlrc5/NLRC5, Tap1/TAP1, and Tap2/TAP2 were measured by quantitative RT-PCR. Data represent the mean ± SEM of the fold change (FC) relative to the siCTRL DMSO from 3 independent experiments. Statistical significance was determined by ordinary one-way ANOVA (*p < 0.05; **p < 0.01; ***p < 0.001; ****p < 0.0001).

**Supplementary Figure 5.**
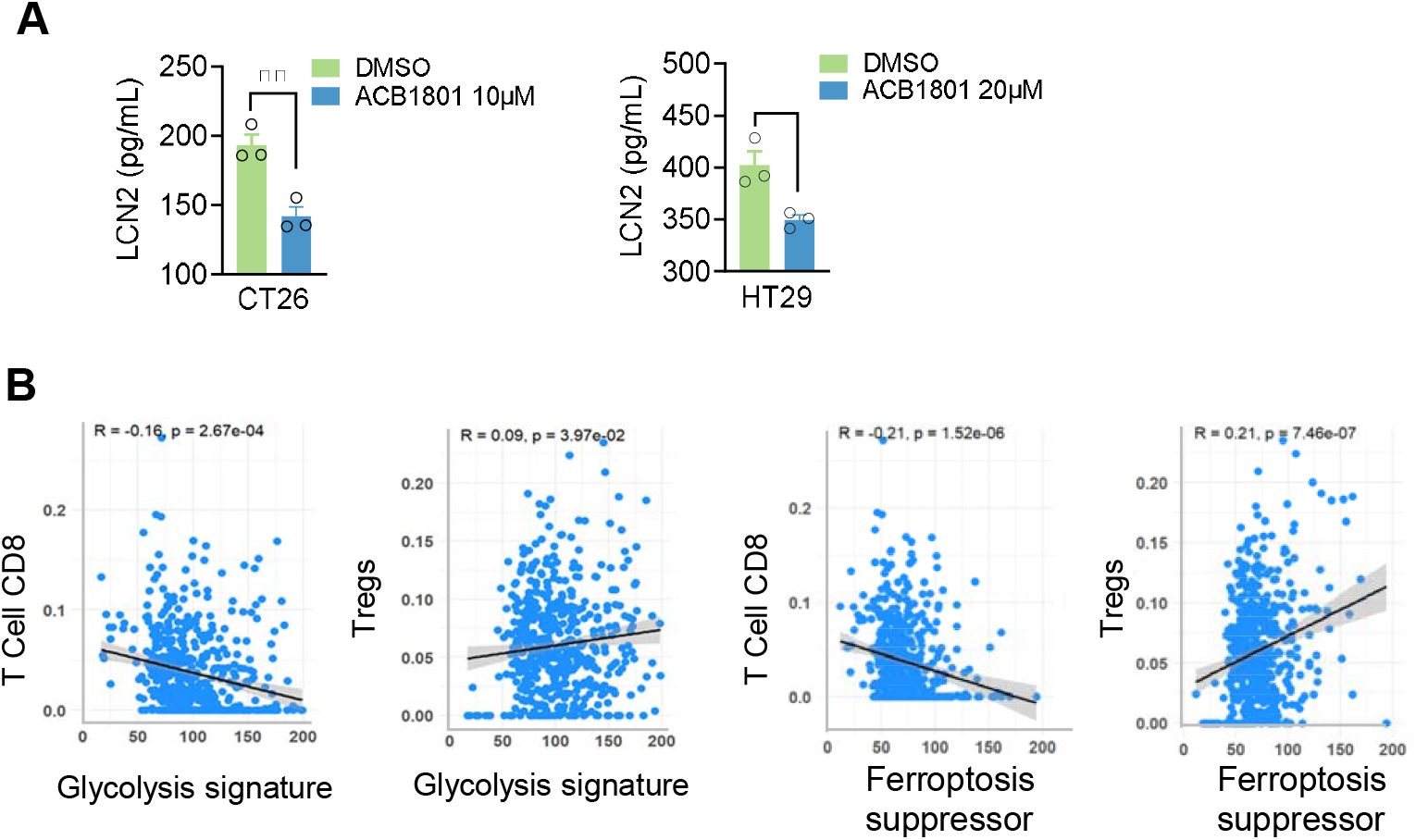
**(A)** ACB1801 reduces the secretion of LCN2 in CT26 and HT29 cells. The bar graphs show LCN2 protein levels in the supernatants of CT26 cells treated with DMSO or 10⍰μM ACB1801, and HT29 cells treated with DMSO or 20⍰μM ACB1801. ACB1801 treatment significantly decreases LCN2 secretion in both cell lines. Data are presented as mean ± SEM from 3 independent experiments. Statistical significance was determined by an unpaired t-test (*p < 0.05, **p < 0.01). **(B)** Association of glycolysis and ferroptosis signatures with T cell subsets in TCGA colorectal cancer (CRC) patients. The scatter plots show correlations between glycolysis signature scores or ferroptosis suppressor scores and the estimated fractions of CD8 T cells or regulatory T cells (Tregs) in CRC tumors from TCGA. A negative correlation is observed between glycolysis or ferroptosis suppressor signatures and CD8 T cell marker, whereas a positive correlation is seen with Treg marker, as indicated by the Pearson correlation coefficients (R) and p-values shown on each plot.

