## Supplementary figures and images for "ACB1801 enhances tumor immunogenicity by targeting glycolysis/ferroptosis vulnerability and activating STAT1-signaling to overcome anti-PD-1 resistance in MSS colorectal cancer"

### Supp Fig 1

#### Supplementary Figure 1

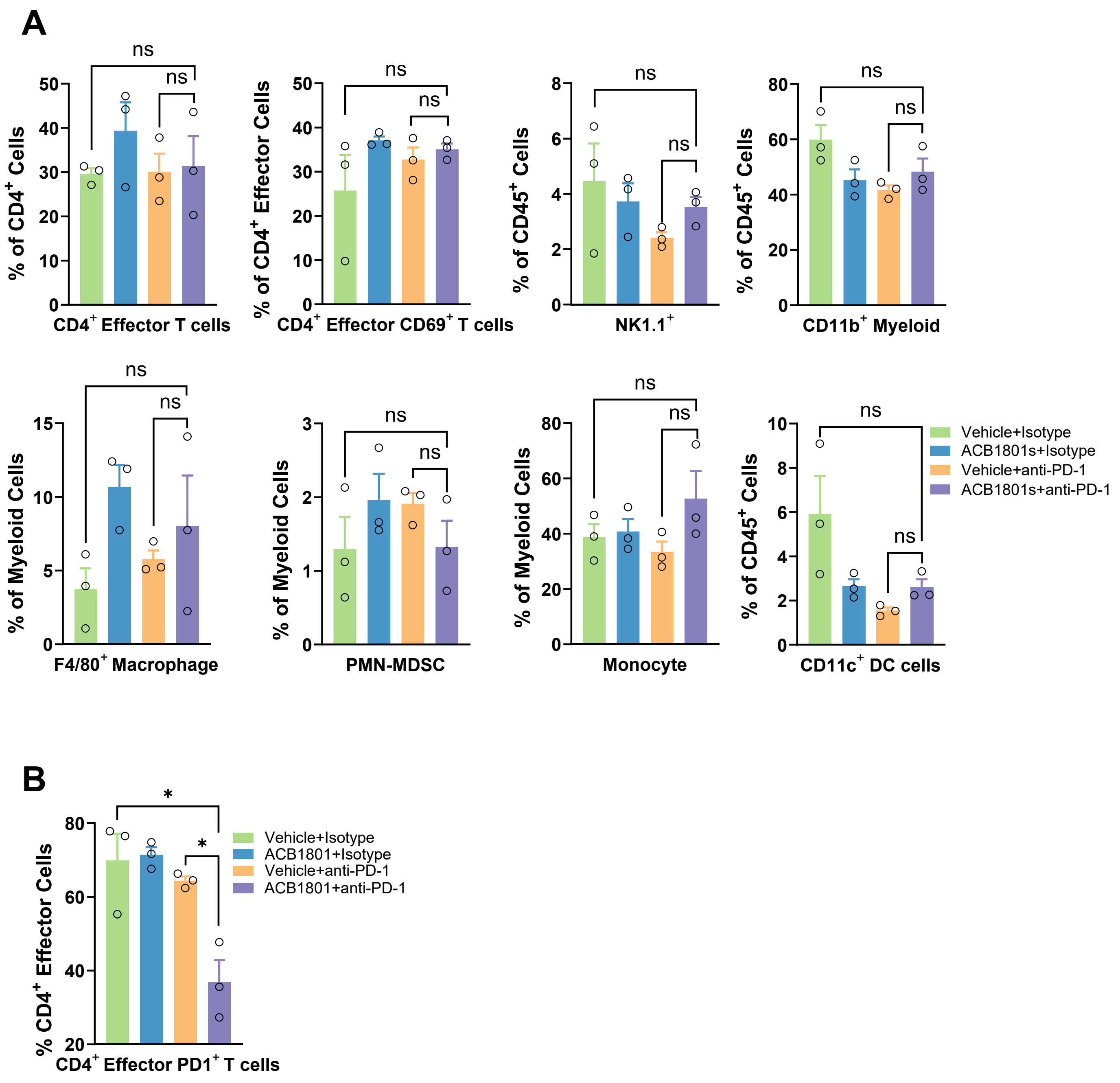

### Supp Fig 2

#### Supplementary Figure 2

**A**

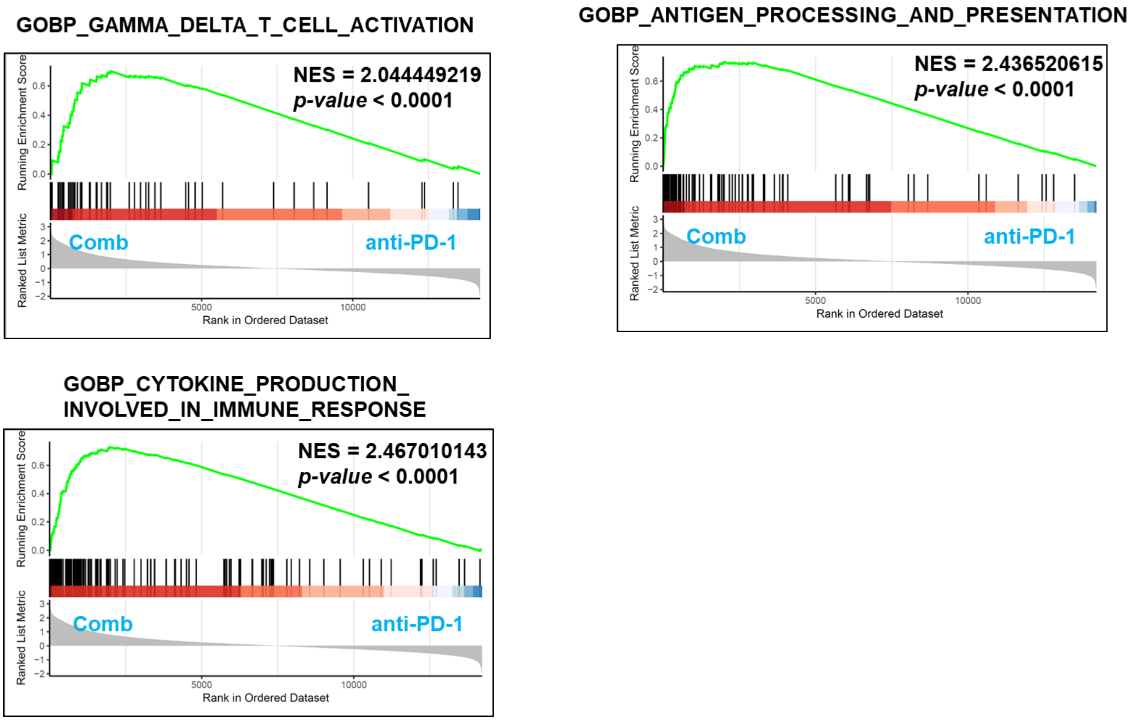


**B**

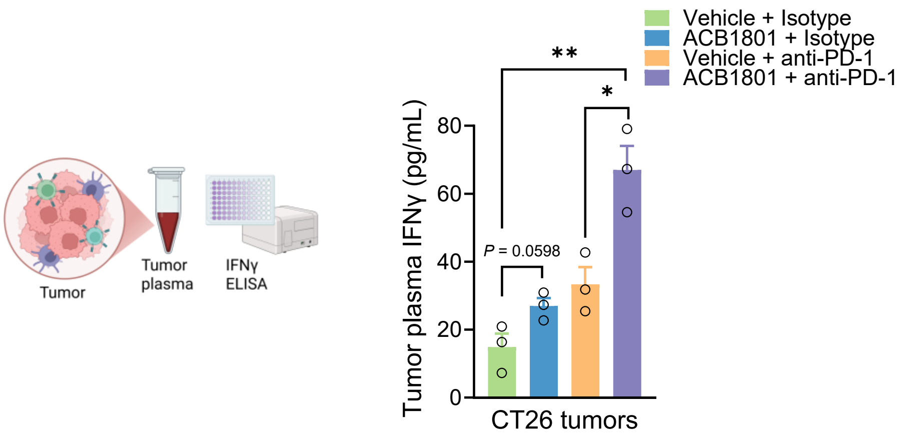

### Supp Fig 3

#### Supplementary Figure 3

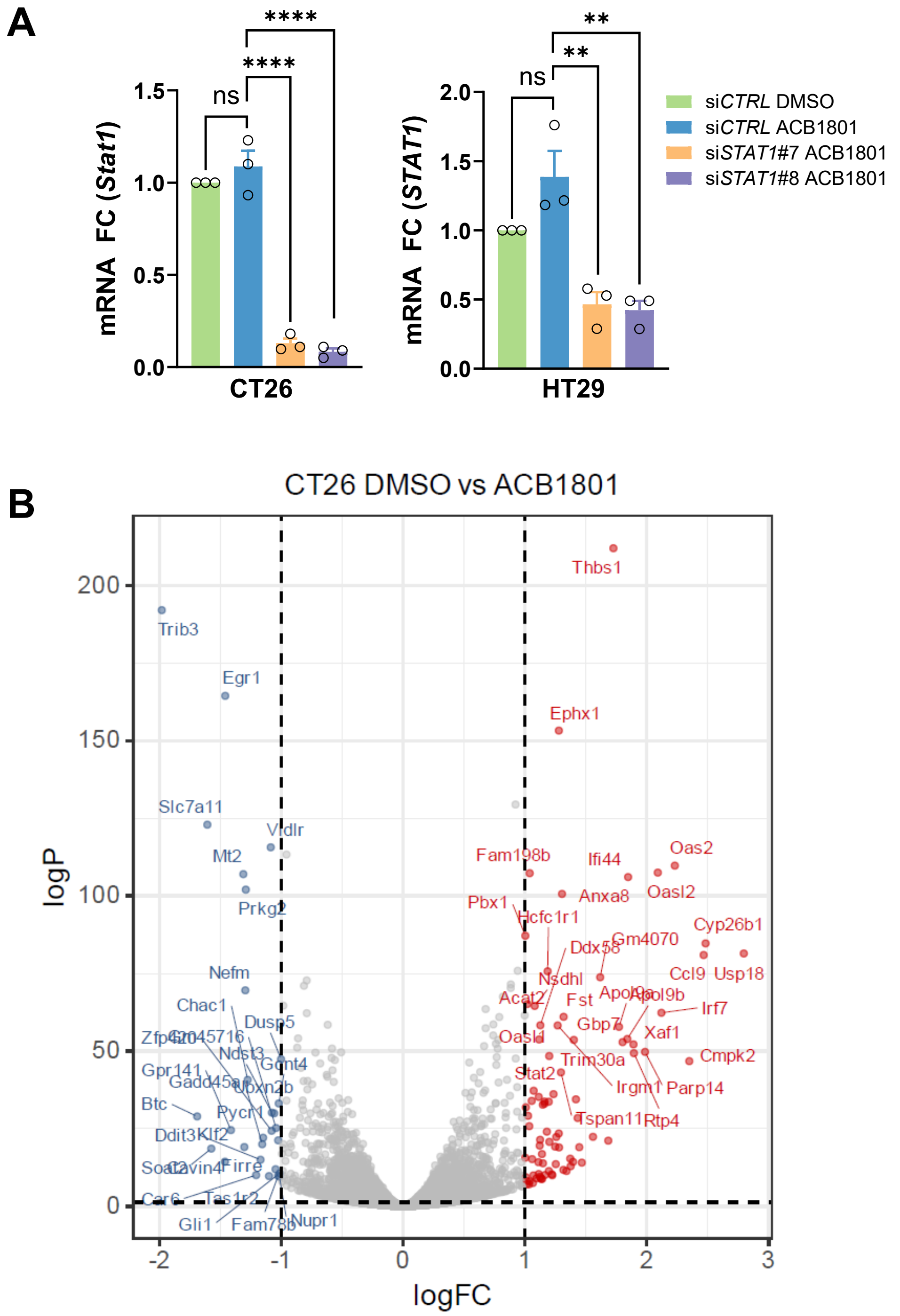

### Supp Fig 4

#### Supplementary Figure 4

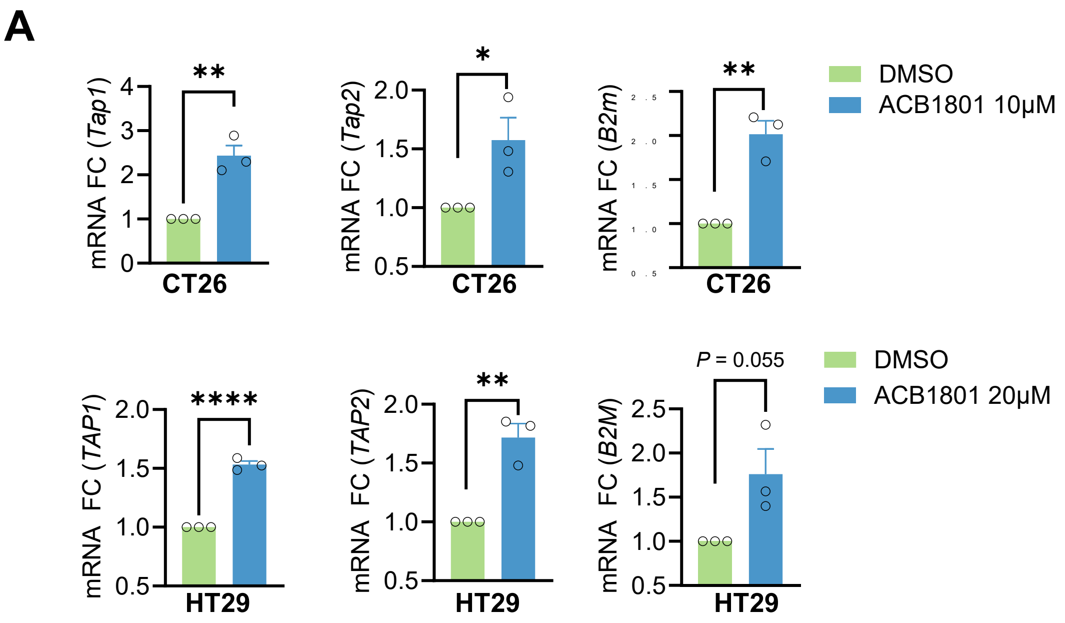


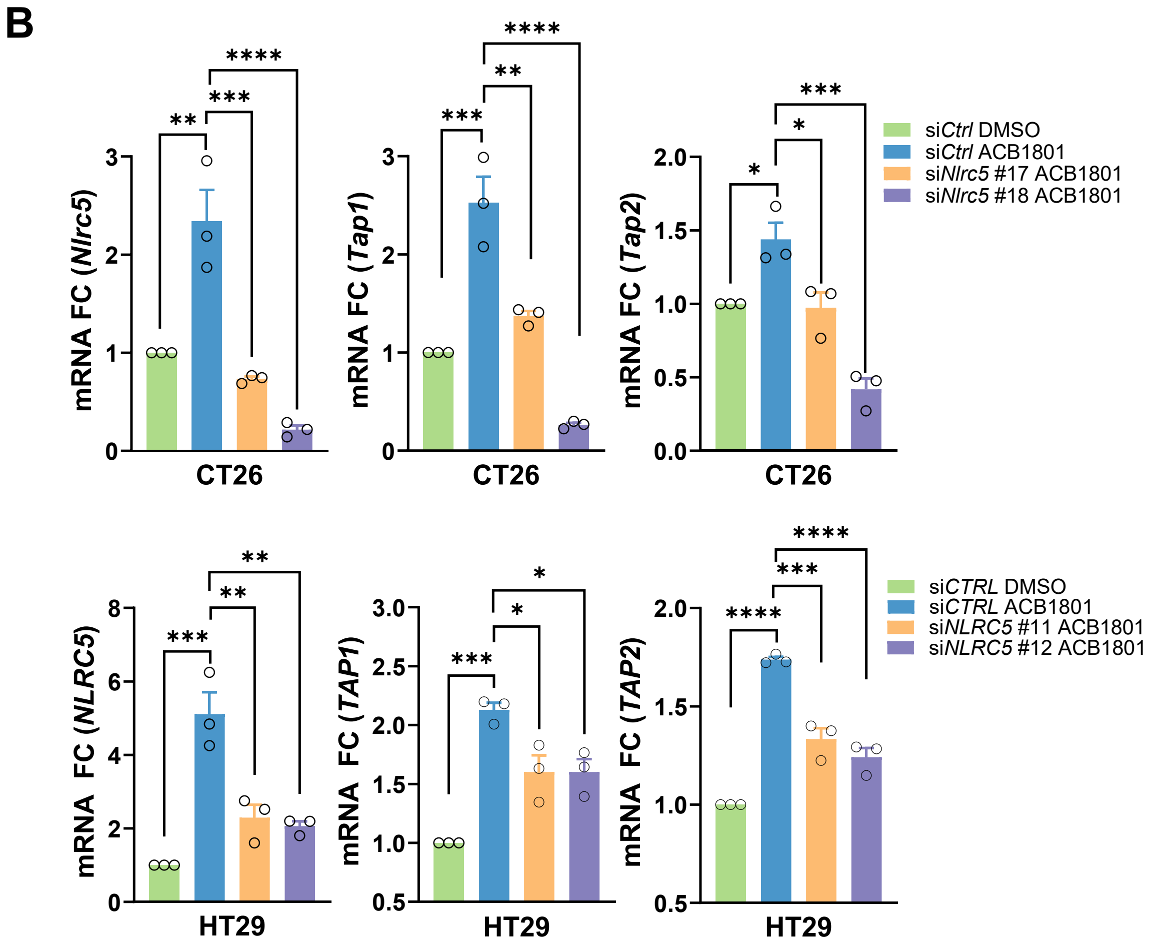

### Supp Fig 5

#### Supplementary Figure 5

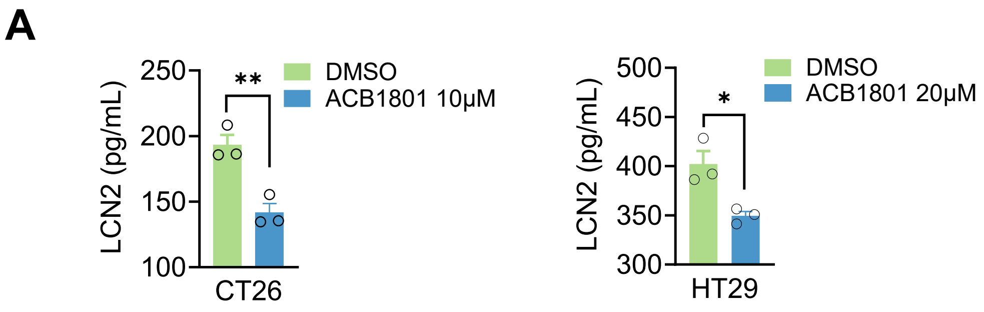


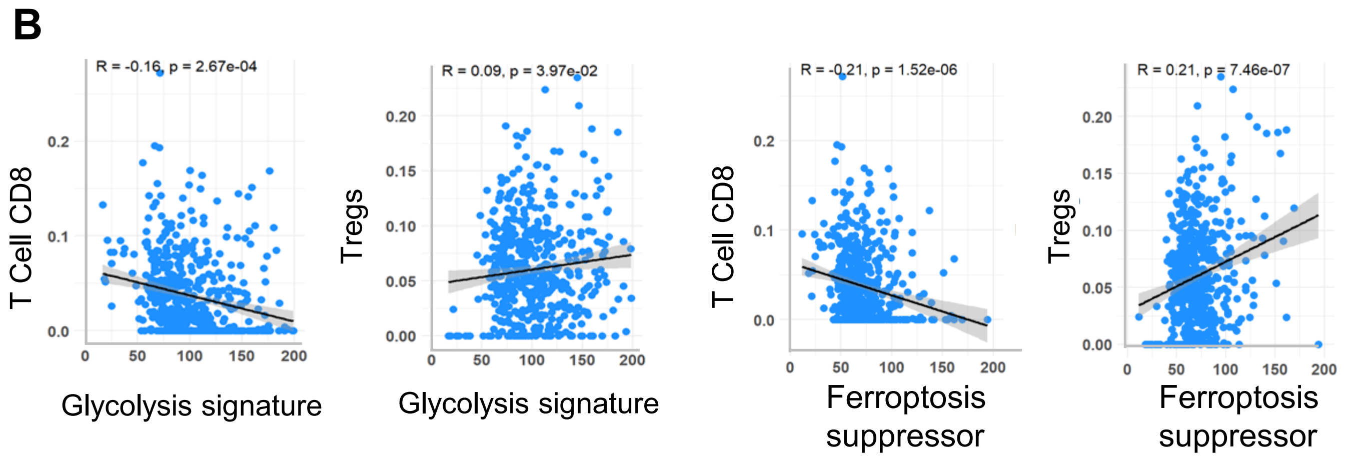
